# Sex differences in autoimmunity could be associated with altered regulatory T cell phenotype and lipoprotein metabolism

**DOI:** 10.1101/760975

**Authors:** George A Robinson, Kirsty E Waddington, Marsilio Adriani, Anna Radziszewska, Hannah Peckham, David. A Isenberg, Yiannis Ioannou, Coziana Ciurtin, Ines Pineda-Torra, Elizabeth C Jury

## Abstract

Male and female immune responses are known to differ resulting in an increased prevalence of autoimmunity in women. Here sex differences in T-cell subset frequency and function during adolescence were examined in healthy donors and patients with the autoimmune disease juvenile (J)SLE; onset of JSLE commonly occurs during puberty suggesting a strong hormonal influence. Healthy adolescent males had increased regulatory T-cell (Treg) frequency, and increased Treg suppressive capacity and IL-4 production compared to healthy adolescent females. The T-helper 2-like profile in male Tregs was associated with increased expression of GATA3 which correlated significantly with elevated Treg plasma membrane glycosphingolipid expression. Differential Treg phenotype was associated with unique serum metabolomic profiles in males compared to female adolescents. Notably, very low density lipoprotein (VLDL) metabolomic signatures correlated positively with activated Tregs in males but with resting Tregs in females. Consistently, only VLDL isolated from male serum was able to induce increased Treg IL-4 production and glycosphingolipid expression following in cultured cells. Remarkably, gender differences in Treg frequency, phenotype and function and serum metabolomic profiles were lost in adolescents with JSLE. This work provides evidence that a combination of pubertal development, immune cell defects and dyslipidemia may contribute to JSLE pathogenesis.

## INTRODUCTION

Males and females differ in their immune response to foreign and self-antigens and consequently they differ in their risk of infection and prevalence of autoimmune diseases; males are generally more susceptible to infections than females, and females represent ~80% of all patients with autoimmunity [1]. It is known that fundamental differences exist in the frequency and activity of T-cell subsets by gender across multiple ethnicities [2–4]. Notably, some gender differences in adaptive immune responses are present throughout life, while others are manifested following the onset of puberty and prior to reproductive senescence implicating both genetic and hormonal influences [5]. Counter to the prevalence of autoimmunity in females, risk of developing cardiovascular disease and relative risk of hypercholesterolaemia is lower in pre-menopausal healthy females compared to males. This is largely due to reduced pro-atherogenic low density lipoproteins (LDL) and increased anti-atherogenic high density lipoproteins (HDL) in females. Following menopause, female LDL increases to a higher level to males, thus reducing their cardiovascular disease protection. HDL remains higher in females at all ages [10–12]. The interaction between serum lipids, sex hormones and immune cell phenotype in males and females remains largely unknown.

Systemic lupus erythematosus (SLE) is a complex autoimmune disorder characterized by loss of immune cell regulation and the presence of autoantibodies against multiple nuclear and non-nuclear antigens, resulting in chronic inflammation, multiple organ damage. Hormones have been implicated in the etiology of SLE due to the sexual dysmorphism observed in SLE where the female to male ratio is 9:1 [1]. Between 15 and 20% of all patients with SLE have juvenile-onset disease (JSLE) which is characterised by a more severe disease phenotype [6][7]. Importantly, cardiovascular disease (CVD) is recognised as a serious long term complication for all patients with SLE [8]. Interplay between traditional cardiovascular risk factors (such as dyslipidemia) and risk factors associated with continuing SLE disease (including hormonal influences), are not understood.

Regulatory T cells (Tregs) have been implicated in SLE due to their suppressive capabilities in the immune system. In SLE, disproportionate Th17-to-Treg ratios have been reported in several studies resulting in a more pro-inflammatory phenotype [13–15]. In health, Treg frequencies are shown to be lower in females following puberty suggesting a plausible reason for the gender bias of autoimmune disease [16]. No difference in Treg frequencies were reported in Australian infants from birth to one year of age [17]. Controversially it has been found that physiological levels of estrogen can induce the expansion of Treg cells [18] and Treg numbers are increased during the menstrual cycle when estrogen levels are highest before ovulation [19]. Foxp3 is a critical protein responsible for the development and function of Treg cells. The gene for Foxp3 is found on the X-chromosome [20] providing a link between sex and Treg development.

Understanding the relationship between sex and Tregs is important for investigation into the aetiology of autoimmune diseases such as SLE. Here the phenotype and function of Tregs in a young cohort of healthy post-pubertal males and females was assessed. Male Tregs had a higher frequency and increased suppressive capacity compared to females and this was associated with an increased expression of IL-4, GATA3, GLUT-1 (glucose transporter-1) and plasma membrane glycosphingolipids. Serum lipids, particularly very low density lipoproteins (VLDL), were associated with the differential Treg phenotype seen between healthy males and females. These sex differences in Tregs and serum lipoprotein profile were lost in JSLE where female patients develop towards a more male Treg phenotype. This could open a new area of research into the breakdown of hormone signalling in JSLE pathogenesis.

## RESULTS

### Young adolescent healthy males have increased Treg frequency and suppressive capacity compared to females

The increased prevalence of autoimmunity in females has been associated with reduced Treg frequencies compared to males [21]. In healthy adolescent males, Treg (CD4^+^CD25^+^CD127^−^, Supplementary Figure 1 for gating strategy) frequency and absolute number were significantly elevated and responder T cells (Tresp, CD4^+^CD25^−^ CD127^+^) were significantly reduced compared to healthy female donors (matched for age, ethnicity and pubertal state) (Figure 1A-D and Supplemental Table 1 for demographic information of healthy donors), confirming previous findings [1, 16]. No significant differences in the frequency of other CD4^+^ and CD8^+^ T-cell subsets were seen (Figure 1A).

**Figure 1:**
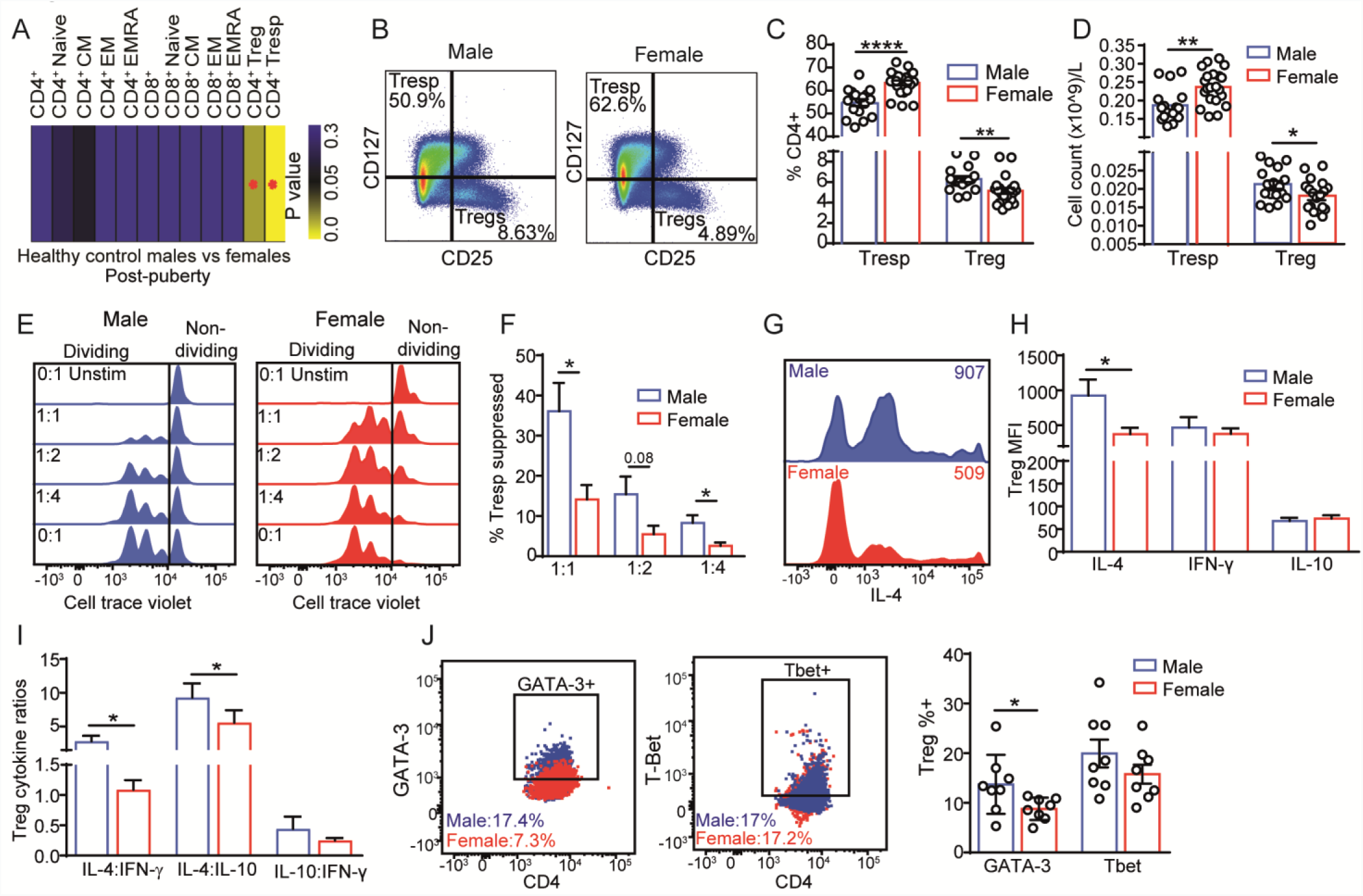
Sex differences in Treg frequency and suppressive capacity. *Ex-vivo* peripheral blood mononuclear cells (PBMCs) from 39 healthy adolescent donors (22 female and 17 male) matched for age, ethnicity and pubertal stage (all puberty tanner stage 4-5) were stained with fluorochrome-labelled antibodies to detect T-cell subset frequencies by flow cytometry (gating strategy Supplementary Figure 1). **(A)** Heatmap of p values comparing T-cell subset immunophenotyping in males vs females. Star=significant p value following 10% false discovery rate adjustment for multiple comparisons (Benjamini, Krieger and Yekutieli approach). **(B)** Representative plots showing responder (Tresp) and regulatory (Treg) T-cell frequencies in post-pubertal males and females. **(C)** Cumulative data of Treg and Tresp cell frequencies and **(D)** absolute counts from 22 post-pubertal females and 17 males. Absolute counts were calculated by multiplying the percentage of Tregs in the total lymphocyte count by the average University College Hospital (UCLH) healthy donor lymphocyte count (x10^9 cells/L). Samples were analysed using Fortessa X20 flow cytometer and Flowjo software. Mean+SE, t-test, *=p<0.05, **=p<0.01, ***=p<0.001, ****=p<0.0001. **(E-F)** Treg mediated suppression of activated Tresp cells. Treg, Tresp and monocytes were isolated from healthy donors (5 male and 5 female). Tresp cells were stained with cell trace violet (CTV) and activated using soluble anti-CD3/28. Monocytes were added to the culture (50,000 cells/well) and Tregs at Treg:Tresp ratios of 1:1, 1:2, 1:4 and 0:1. After 72hrs, proliferation (CTV) was measured using Fortessa X20 flow cytometer and Flowjo software (Supplemental Figure 2 and 3 for sorting and gating strategies). **(E)** Representative plots showing proliferation of Tresp cells at varying Treg:Tresp ratios in males (left) or females (right). Each leftward peak represents a round of proliferation. **(F)** Cumulative data showing suppression capacity of Tregs at varying Treg:Tresp ratios in males compared to females. This was calculated using the fold change of Tresp % proliferation with Tregs (1:1, 1:2, 1:4) compared to Tresp without Tregs (0:1). Mean+SE, t-test, *=p<0.05. **(G-I)** *Ex vivo* T-cells from 8 male and 12 female healthy donors were surface stained for Treg markers and intracellularly for cytokines IL-4, IFN-γ and IL-10 after 4hr stimulation with PMA/ionomycin/golgi plug. **(G)** Representative plots displaying IL-4 MFI expression in males and female Tregs. **(H)** cumulative data showing Treg IL-4, IFNγ and IL10 expression and **(I)** cytokine expression ratios in Tregs from male and female healthy donors. **(J)** Representative plots from *ex vivo* Tregs stained intracellularly for GATA-3 or Tbet and cumulative data showing GATA-3 and Tbet expression in Tregs from 8 healthy males and 8 females. Samples were analysed using Fortessa X20 flow cytometer and Flowjo software. Mean+SE, t-test, *=p<0.05.

In addition to increased number, Tregs from adolescent healthy males had a significantly greater capacity to suppress activated Tresps compared to female Tregs even at higher dilutions, suggesting increased Treg functionality in males (Figure 1E and F and Supplemental Figure 2 and 3 for cell sort and suppression assay gating strategy). This was accompanied by significantly increased IL-4 production and increased IL-4:IFN-γ and IL-4:IL10 ratios by Tregs from healthy male compared to female donors detected by intracellular staining. No differences were observed in Treg IL-10 and IFN-γ production or IL-10:IFN-γ ratio (Figure 1G-I) or IL-4, IL-10 and IFN-γ production by Tresp (Supplemental Figure 4) between males and females. In support of increased IL-4 production by male Tregs, GATA-3 (the transcription factor associated with an IL-4/Th2 phenotype [22]) was also significantly increased in healthy male compared to female Tregs whilst no difference was seen in T-bet expression (Figure 1J).

### Tregs from adolescent males have a more metabolically active phenotype characterised by increased plasma membrane glycosphingolipids and Glut-1 expression that correlates with a Th2 phenotype

Further analysis of male and female Treg phenotype revealed a more metabolically activated profile in adolescent males characterised by increased GLUT-1 and lipid raft expression (assessed using cholera-toxin-B, CTB to measure plasma membrane glycosphingolipids (GSL) [23]), although expression of activation markers CD69 and PD-1 remained unchanged (Figure 2A-C).

**Figure 2:**
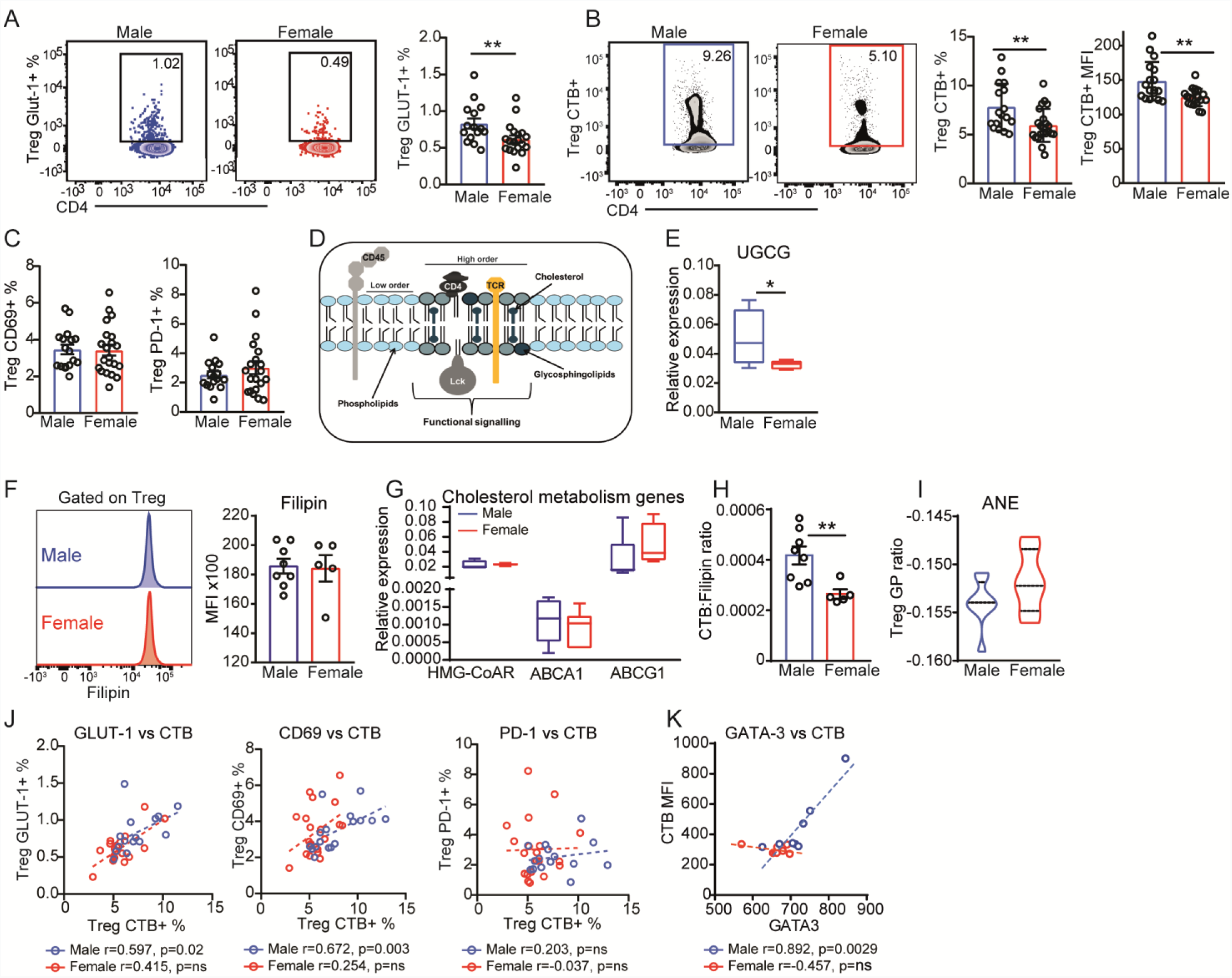
Th2 phenotype of Tregs in males related to lipid metabolism. Assessment of metabolic and functional profile of male and female Tregs. Healthy donor PBMCs from 22 females and 17 males were stained with fluorochrome labelled antibodies (CD4, CD25 and CD127) to detect Treg cells and functional (CD69 and PD-1) and metabolic markers GLUT-1 and plasma membrane lipids (cholera toxin-B, CTB assessing glycosphingolipids, and filipin binding to cholesterol). **(A)** Representative plots (left) and cumulative expression levels (right) of GLUT-1 in Tregs. **(B)** Representative plots (left) and cumulative expression levels (right) of CTB expression. **(C)** cumulative expression levels of Treg CD69 (left) and PD-1 (right). **(D)** T-cell lipid raft and membrane order model highlighting key components cholesterol and glycosphingolipids (GSL) as well as the T-cell receptor (TCR) and signalling proteins. **(E)** RNA isolated from FACS-Aria sorted Treg (CD4+CD25hiCD127-) cells from 4 healthy males and 4 females and analysed by qPCR for gene expression of UGCG (UDP-Glucose Ceramide Glucosyltransferase). Relative gene expression compared to cyclophilin. **(F)** Representative plots (left) and expression levels (right) of cholesterol (filipin) expression in Tregs from 8 males and 5 females. **(G)** RNA isolated from FACS-Aria sorted Treg (CD4+CD25hiCD127) cells from 4 healthy males and 4 females and analysed by qPCR for gene expression associated with cholesterol metabolism: ATP-binding cassette transporter A1 (ABCA1), ATP-binding cassette transporter G1 (ABCG1), 3-hydroxy-3-methyl-glutaryl-coenzyme A reductase (HMGCR). Relative gene expression compared to cyclophilin. **(H)** Ratio of GSL:cholesterol in Tregs from 8 males and 5 females. **(I)** Live cells were stained with fluorochrome labelled antibodies (CD4, CD25 and CD127) to detect Treg cells and subsequently stained with di-4-ANEPPDHQ (ANE) to detect membrane order (using calculated generalised polarisation (GP) ratios) by flow cytometry as previously described [25]. **(A-I)** Mean+SE, t-test, *=p<0.05, **=p<0.01. **(J)** Expression of GSL correlated with GLUT-1, CD69 and PD-1 in matched Tregs from 22 females and 17 males and **(K)** GATA-3, measured intracellularly, in matched Tregs from 8 males and 8 females. P-value and correlation coefficients are displayed from the Pearson correlations.

Our previous work shows that T-cell activation and polarisation is influenced by changes in the plasma membrane lipid profile (Figure 2D [24, 25]). The increase in GSL expression in male Tregs was matched by elevated UDP-Glucose Ceramide Glucosyltransferase (UGCG) gene expression, the rate limiting enzyme in the GSL synthesis pathway [26] (Figure 2E). No differences were seen in Treg cholesterol expression (Figure 2F), or cholesterol biosynthesis (HMG-CoAR) and cholesterol efflux (ABCA1 and ABCG1) gene expression between males and females (Figure 2G). However, the GSL:cholesterol ratio was significantly elevated (Figure 2H) and the plasma membrane lipid order (measured by the phase sensitive probe ANE) (Figure 2I) was reduced suggesting that the plasma membrane of male Tregs was more fluid (less ordered) compared to female Tregs. Finally, a significant positive correlation was observed between male Treg plasma membrane GSL levels and GLUT-1, CD69 and GATA-3 expression (Figure 2J and K) implying that differences in Treg function between males and females may be related to a differential lipid metabolism.

### Healthy adolescent males and females have unique serum metabolomic profiles associated with differential Treg phenotypes

The onset of puberty is known to alter lipid metabolism [27–31] and these differences could contribute to the changes in plasma membrane lipid profiles and function seen in male compared to female Tregs [32]. Following an in depth analysis of serum lipid metabolites using an established nuclear magnetic resonance (NMR) spectroscopy platform (https://nightingalehealth.com/) (Supplemental Table 2), 42 metabolites were significantly associated with adolescent post-pubertal males and 24 with females (Figure 3A, Supplemental Table 3). Notably, a significant increase in VLDL particle subsets (including size and lipid composition) was identified in males, and a significant increase in HDL-particles was seen in females (Figure 3A, Supplemental Table 3). This was supported by an increase in Apolipoprotein (Apo)-B in males and ApoA1 in females. Furthermore, there was a significant increase in serum polyunsaturated fatty acids (PUFA’s) in adolescent females and increased saturated fatty acids SFA’s and serum triglycerides (TG) in adolescent males. Unbiased hierarchical clustering using the significantly altered metabolites between males and females was able to differentiate two clusters. Cluster-1 contained 79% of the males with VLDL measures as the predominant metabolites. Cluster-2 contained 76% of the females with HDL measures as the predominant metabolites (Figure 3B). Principle component analysis (PCA) was similarly effective in clustering males from females using the significantly altered metabolites between males and females (Figure 3C).

**Figure 3:**
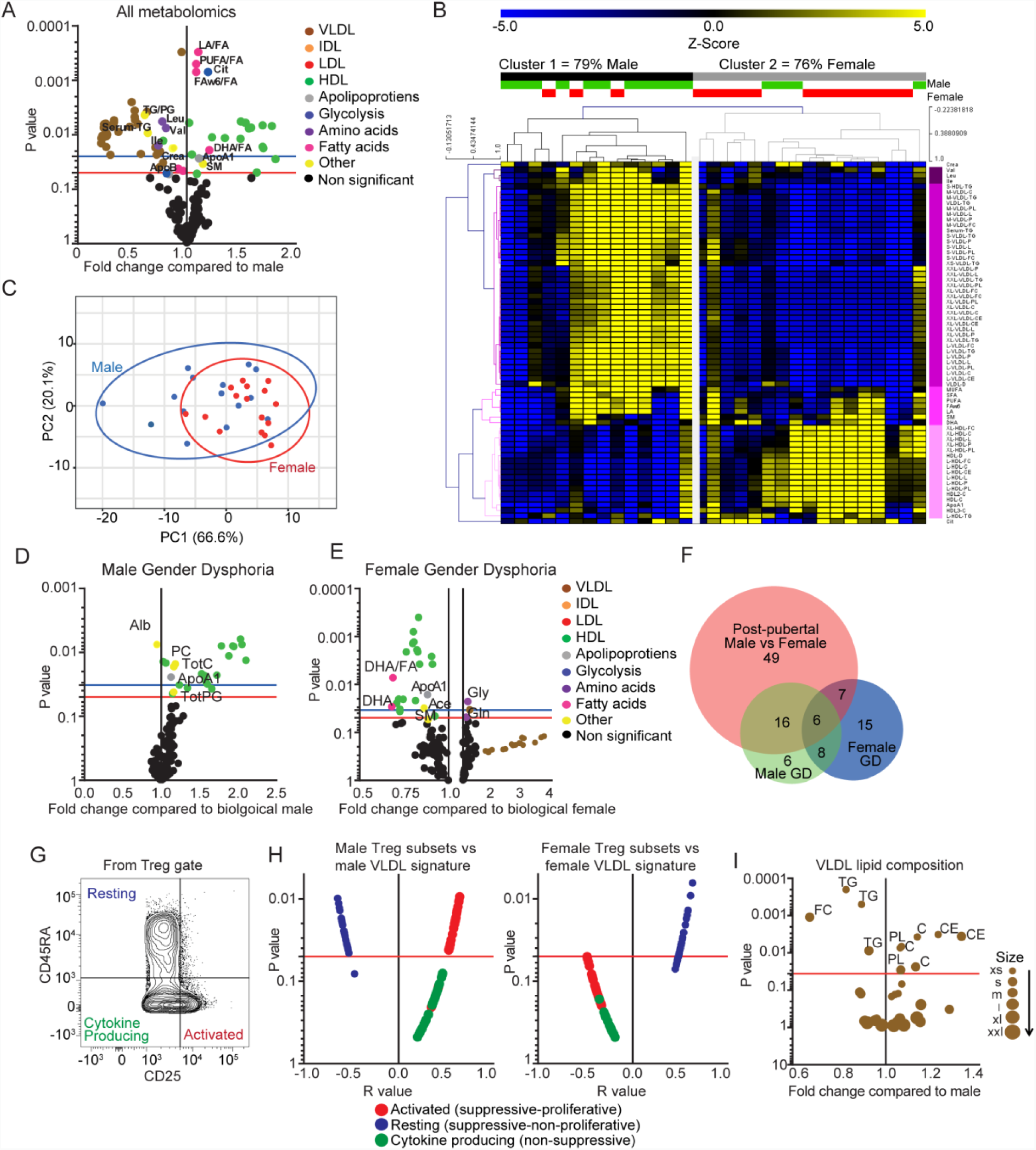
Young males and females have an altered serum lipid profile associated with Treg phenotype. Metabolites were measured in male or female serum (Supplementary Table 2). **(A)** Volcano plot displaying fold change of all metabolites (from male expression levels) and Log10 p values from multiple unpaired t-tests comparing healthy males (n=15) and females (n=17). **(B)** Heat map displaying the z-score converted individual measurements of the male and female metabolic signatures (Supplemental Table 3) from 32 individual healthy controls. Unbiased hierarchical clustering (Pearsons) was performed using MeV software (males=green, females=red). **(C)** Principle component analysis performed using the significant metabolites between post-pubertal males (n=15, blue) and females (n=17, red) identified in (A) (Supplemental Table 3). **(D-E)** Volcano plot showing fold change of gender dysphoria (GD) individuals (n=10) from **(D)** respective male (XY, n=15) or **(E)** respective female (XX, n=17) expression levels and Log10 p values. **(A, D and E)** Blue line=p=0.05, Red line=adjusted p value threshold following 10% false discovery rate adjustment for multiple comparisons (Benjamini, Krieger and Yekutieli approach). Coloured dots represent different metabolite groups. Abbreviations: VL/I/L/HDL (very low/intermediate/low/high density lipoproteins), C (cholesterol), CE (cholesterol ester), FC (free cholesterol), TG (triglycerides), PL (phospholipids), Apo (apolipoprotein), TG (triglycerides), FA (fatty acids), PG (Phosphoglyceride), PC (Phosphatidylcholine), SM (Sphingomyelins), DHA (Docosahexaenoic acid), LA (Linoleic acid), FAw6 (Omega-6 fatty acids), PUFA (Polyunsaturated fatty acids), MUFA (Monounsaturated fatty acids), SFA (Saturated fatty acids). **(F)** Venn diagram displaying the number of significantly different metabolites (Supplemental Table 4) that overlap between different gender group comparisons. GD; Gender dysphoria. **(G)** Flow cytometry PBMC gating strategy for Treg functional subsets described by Miyara et.al [33]. Healthy donor PBMCs were stained with fluorochrome-labelled antibodies (CD4, CD25, CD127 & CD45RA). **(H)** Volcano plot showing the correlation (r value) between the male (left) or female (right) VLDL signature (Supplemental Table 3) and Treg functional subsets and Log10 p values. Red line=p=0.05, coloured dots represent different Treg functional subsets. **(I)** Volcano plot showing VLDL percentage lipid compositions between healthy males and females. Data is displayed as fold change from male expression levels and Log10 p values of healthy males (n=15) and females (n=17). Red line=adjusted p value following 10% false discovery rate adjustment for multiple comparisons. Dot size represents different sized VLDL particles.

To investigate the hypothesis that sex hormones can influence lipid metabolism, we explored the serum lipid profile of males and females with gender dysphoria (GD). Biological males (XY) with inhibited testosterone and supplemented estrogen had an increase in HDL metabolites including ApoA1 (Figure 3D) resembling the profile of young post-pubertal females. In contrast, biological females (XX) with estrogen inhibition and testosterone supplementation had decreased HDL, ApoA1 and the PUFA docosahexaenoic acid (DHA) and up to 4-fold increased VLDL expression (Figure 3E) resembling the profile of young post-pubertal males. From the metabolites significantly altered between males and females, proportional Venn analysis identified 16 metabolites that overlap with the male GD analysis and 7 that overlap with the female GD analysis (Figure 3F, Supplemental Table 4). A further 8 metabolites were common between male and female GD donors and 6 metabolites overlapped with all the analysis comparisons. Thus supporting that hormones play a role in serum lipid metabolism.

When the sex specific metabolic signatures were correlated with male and female Treg frequency, no relationships were identified (Supplemental Figure 5A). However, when functional Treg subsets were considered (determined using the gating strategy described by Miyara et.al [33], Figure 3G) differential correlations were detected. In males, VLDL correlated positively with activated Tregs (CD25^++^CD45RA, proliferative and suppressive) and negatively with resting Tregs (CD25^+^CD45RA^+^, non-proliferating and suppressive) (Figure 3H). In females, VLDL had a significant positive correlation with resting Tregs. A more detailed analysis of VLDL particles revealed that male small and medium sized VLDL contained significantly more triglycerides and free cholesterol whereas VLDL from females contained more cholesterol, cholesterol esters and phospholipids (Figure 3I).

No associations were identified between HDL and Treg phenotype (Supplemental Figure 5B). Thus, healthy adolescent males and females had Tregs with a differential membrane lipid and metabolic profile and significantly different serum lipid signatures that correlated with different Treg functional phenotypes.

### Differences in VLDL composition between males and females drive differences in Treg phenotype in vitro

The results so far show that the composition of the VLDL particles in adolescent males and females (post puberty) are different. To test the functional impact of these differences in VLDL composition on Tregs, healthy human PBMC’s from anonymised leukocyte cones were rested in serum free medium and then stimulated with VLDL isolated from healthy male and female serum samples (Figure 4A for experimental plan). Culture with male VLDL recapitulated the ex vivo male Treg phenotype, characterised by increased IL-4 production and increased GSL expression compared to PBMCs cultured with female VLDL (Fig 4B-D). Conversely, female VLDL significantly increased Treg phospholipid expression compared to male VLDL (Figure 4E). No significant differences were detected for Treg cholesterol (Figure 4F), IL-10 or IFN-γ production as we described previously (Figure 1H). Thus, healthy Treg phenotype can be differentially altered by VLDL isolated from males or females.

**Figure 4:**
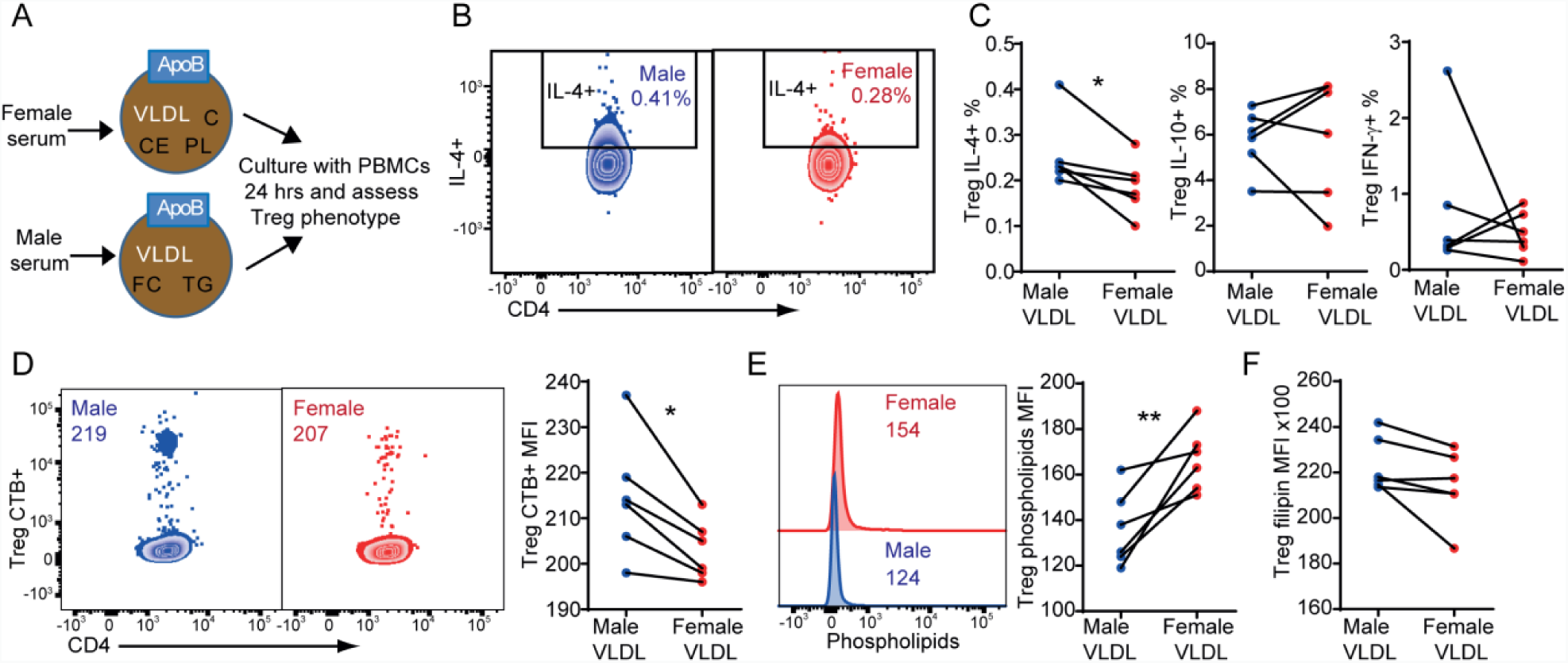
Isolated VLDL from male and female serum induces different phenotypic characteristics in Tregs in vitro. Association of male or female VLDL with Treg phenotype and function *in vitro*. **(A)** Experimental design for the isolation of VLDL from male or female serum and culture with PBMC’s. Healthy donor PBMC’s cells from 6 leukocyte cones (gender unknown but rested in serum free medium before culture) were stimulated with VLDL isolated from male or female serum (50ug/ul) for 24hrs and then stained with fluorochrome labelled antibodies (CD4, CD25 and CD127) to detect Tregs. Cells were stained intracellularly for cytokines IL-4, IL-10 and IFN-γ following subsequent 4hr stimulation with PMA/ionomycin/golgi plug. C, cholesterol; CE, cholesterol esters; FC, free cholesterol; TG, triglycerides; PL, phospholipids. **(B)** Representative plots of IL-4 and **(C)** cumulative expression levels of cytokines IL-4, IL-10 and IFN-γ. Cells were separately stained with **(D)** CTB to detect PM GSL, **(E)** lipidtox detection kit to detect phospholipids and **(F)** filipin to detect cholesterol. Mean. Paired t-tests, *=p<0.05, **=p<0.01

### Sex specific differences in metabolism and Treg phenotype are lost in JSLE

The reduced number and suppressive capacity of Tregs in adolescent females could play a role in the increased susceptibility of females to autoimmune diseases. Juvenile-onset (J)SLE has a substantial female sex bias, with a 4.5:1 female:male ratio [34]. The T cell phenotype was examined in male and female patients with JSLE who were age matched to both each other and to the adolescent healthy controls (Supplemental Table 5). Strikingly, the sex-associated differences identified between Tregs from healthy donor males and females were lost in patients with JSLE; this was true for Treg frequency, lipid profile, and IL-4 production (Figure 5A-D). Significant differences in T-cell phenotype, including Tregs, were identified in JSLE females when compared to healthy females but not for JSLE males when compared to healthy males (Supplemental Figure 6).

**Figure 5:**
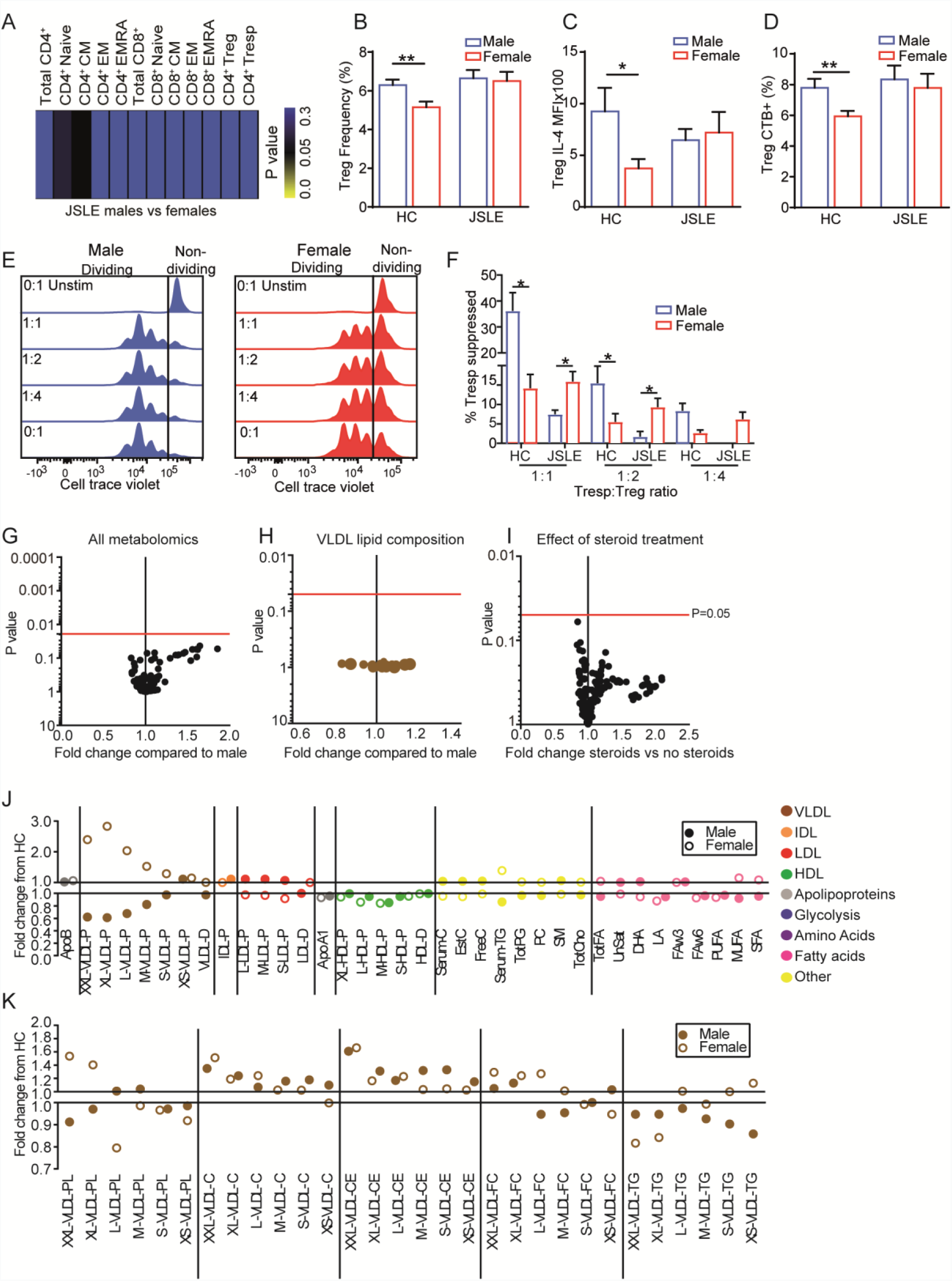
Sex differences in metabolism and Treg phenotype are lost in JSLE. **(A)** Heatmap of p values comparing T-cell subset immunophenotyping (gating strategy in Supplemental Figure 1) of 39 age and ethnically matched JSLE patients (23 female and 12 male). Unpaired t-test. **(B)** JSLE PBMCs stained with fluorochrome-labelled antibodies (CD4, CD25, CD127) to detect Treg cell frequency measured in 22 female and 17 male healthy donors and 23 female and 12 male JSLE patients. **(C)** Ex vivo T-cells from 8 male and 12 female healthy donors and 6 male and 15 female JSLE patients surface stained for Treg markers and intracellularly for cytokine IL-4 after 4hr stimulation with PMA/ionomycin/golgi plug. **(D)** PBMCs as described in (B) were stained with fluorochrome labelled antibodies (CD4, CD25 and CD127) to detect Tregs and CTB (GSL) PM lipid. Mean. t-tests, *=p<0.05, **=p<0.01. **(E-F)** Treg mediated suppression of activated Tresp cells as measured by cell trace violet (CTV) proliferation detection agent) in JSLE patients (6 male and 6 female) and compared to healthy controls as described in Figure 1E and 1F (Supplemental Figure 2 and 3 for gating strategies). **(E)** Representative plots showing the extent of proliferation of Tresp cells at varying Treg:Tresp ratios in JSLE males (left) or JSLE females (right). Each leftward peak represents a round of proliferation. **(F)** Suppression capacity of Tregs at varying Treg:Tresp ratios in males compared to females calculated as described in Figure 1F. Mean+SE, t-test, *=p<0.05. **(G)** Volcano plot showing all metabolomics data (Supplemental Table 2) and **(H)** VLDL compositional differences from 11 JSLE males and 22 JSLE females. Data is displayed as fold change from male expression levels and Log10 p values. **(I)** Volcano plot showing metabolomics data from steroid treated (n=17) and non-treated (n=16) with expression levels displayed as fold change from non-treated and Log10 p values. **(G-I)** Log10 p values corrected for multiple testing by 10% false discovery rate (Benjamini, Krieger and Yekutieli approach). Red line=0.05. Dots represent an individual metabolite. **(J)** Lipoprotein particle concentration and **(K)** lipid measures from healthy/JSLE males (n=15/11) and females (n=17/22). Data is displayed as mean fold change from healthy. Male points are represented as filled dots, female as hollow dots.

Interestingly, the suppressive capacity of the Tregs was increased in JSLE females compared to JSLE males, the opposite to what we found in healthy individuals (Figure 5E-F). This appeared to be due to a loss of suppressive capacity in males with JSLE as opposed to an increase in females with JSLE.

Furthermore, the differential sex-associated serum metabolite profile was also lost in patients with JSLE, including the altered VLDL composition in males vs females (Figure 5G-H). Steroid treatment did not significantly influence metabolite profiles (Figure 5I). We next compared the metabolomic profile of healthy donors and JSLE patients separated by sex (Figure 5J). An increase in VLDL particles was found in SLE females whereas a decrease was observed in males. A subtle decrease in HDL particles was also identified in both JSLE males and females. The lipid composition of VLDL was also altered, characterised by differentially reduced phospholipid (PL), Triglyceride (TG) and free cholesterol (FC) content in male VLDL particles compared to females. Cholesterol (C) and cholesterol esters (CE) increased in all VLDL sizes in both male and female JSLE patients (Figure 5K).

Overall, these findings suggest that an alteration in hormone signalling in JSLE could contribute to changes in lipoprotein profile and composition, leading to a loss of differential male/female blood lipid taxonomy. Changes in lipid metabolism can influence immune cell function by influencing plasma membrane lipids, contributing to increased autoimmunity development and potentially increased cardiovascular risk in females. This hypothesis is summarised in Figure 6.

**Figure 6:**
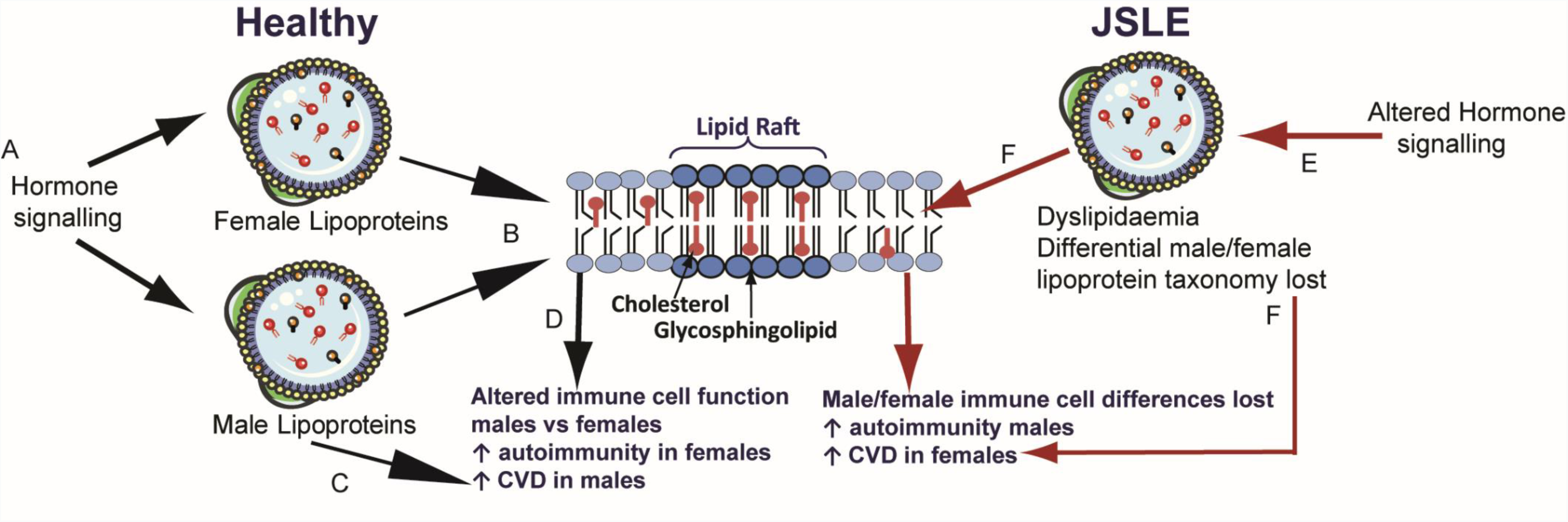
Potential mechanism: How blood lipids influence immune cell function in healthy individuals and JSLE patients. Based on the findings described in this report we hypothesise that in healthy adolescence, (A) hormone signalling via estrogen and/or androgen receptors contribute to changes in blood lipids between males and females including the lipid content and size of VLDL. (B) Altered blood lipid taxonomy can influence immune cell function by changing the lipid content of the plasma membrane. (C) Increased VLDL in males drives increased cardiovascular risk in males; (D) Changes in immune cell activation between males and females can contribute to altered susceptibility to a range of diseases. (E) Potential defects in sex hormone signalling in patients with JSLE lead to a loss of differential male/female blood lipid taxonomy. (F) Blood lipids in JSLE are defective and alter immune cell plasma membrane lipids and immune cell function and increase cardiovascular risk in female JSLE patients.

## Discussion

Here we describe a link between gender, lipid metabolism and Treg phenotype in healthy adolescent donors. Male Tregs had increased peripheral blood frequency, suppressive capacity and IL-4 production compared to Tregs from female donors; this was related to an increased Treg metabolic profile characterised by elevated GLUT-1 expression, plasma membrane lipid expression and plasma membrane fluidity in males compared to females. Healthy adolescent males and females had a distinctive serum metabolic signature including striking differences in VLDL serum lipoprotein particle distribution and lipid content. These signatures were positively correlated with an activated Treg phenotype in males and a resting Treg phenotype in females. In vitro culture with VLDL isolated from males and females was able to appropriately recapitulate the respective male or female Treg phenotype. Finally, sex differences in Treg and serum metabolomic profiles were lost in age matched patients with JSLE, a disease triggered during puberty, suggesting that pathogenesis could be related to dysregulated hormone signalling.

Our findings support previous observations of sexual dimorphism of Treg frequency across different age groups [1, 16], despite evidence that FOXP3 is encoded on the X-chromsome [35]. Tregs from young healthy post-pubertal males were characterised by an increased peripheral blood frequency of Tregs with a Th-2 like phenotype (increased IL-4 production and GATA-3 expression). Th2-like Tregs have a greater viability than other subsets [36] and Treg-specific GATA3 knockout in mice results in spontaneous autoimmune disease development related to the regulation of FOXP3 by GATA3. These GATA3-deficient Tregs developed a pro-inflammatory Th17 phenotype [37, 38]. Differences in T-cell plasma membrane glycosphingolipids and cholesterol (lipid raft components) are known to influence cell function by altering lipid raft structure and membrane fluidity [24, 25, 39, 40]. Cholesterol enrichment in CD4 T-cells results in increased proliferation and Th1 differentiation [41]. In the same study, cholesterol enrichment in Tregs had no effect on function. In support of this, we show that Tregs had no difference in cholesterol expression. However Treg GSL expression and GSL:cholesterol ratio was increased in male compared to female Tregs suggesting the plasma membrane was less ordered and more fluidic in males; this could have functional implications on cell signalling [24, 25]. Furthermore, GLUT-1, CD69 and GATA3 expression correlated positively with GSL levels in males but not in females. IL-4 increases GATA3 expression and Th2 differentiation in T-cells and the levels of GATA-3 are enhanced by concomitant TCR signalling [42]. Increased GSL levels in male Tregs may therefore play a role in driving the Th2 phenotype through increased TCR signalling.

Clinical measures of lipoproteins differ between healthy adult males and pre-menopausal females where males are at an increased risk of developing cardiovascular disease [10–12]. No studies have investigated in depth metabolomics regarding lipoprotein particle size and composition between adolescent healthy males and females. VLDL subsets and their lipid content, saturated and monounsaturated fatty acids were greatly increased in young males and HDL and polyunsaturated fats such as omega fatty acids, DHA and LA were increased in females. It has been previously reported that DHA is lower in males than females, thus supporting these findings [44]. This provides evidence that in adolescence, immediately after puberty, young males and females can already be segregated by metabolic signatures characteristic of their risk of developing cardiovascular disease in later life. It has been shown that the combined oral contraceptive pill (COCP) (combination of an estrogen (ethinylestradiol) and a progestogen (progestin)) increases circulating triglycerides and HDL cholesterol whereas the progestin only oral contraception has little effect on lipids [45]. In the same study the COCP also decreased polyunsaturated fats, together suggesting a complex role for hormones in altering lipid metabolism. Our data supports a role for estrogen in driving the HDL signature in females. The few studies examining lipids and GD however have contradictory observations. One study showed that transgender males develop increased TGs, total cholesterol and LDL-cholesterol as well as decreased HDL-cholesterol. Transgender females in the same study developed decreased total cholesterol and LDL-cholesterol [46]. Another report also identified a decrease in HDL levels in transgender males but found no difference in the lipid profile of transgender females [47]. Finally, a study in mice has shown that during high estrogen periods of the menstrual cycle there is an increased efflux of more efficient, atheroprotective HDL from liver hepatocytes. This is due to the increased co-regulated transcriptional activity of LXR’s by ERα [48]. The hormonal influences we see here may be driven by the same mechanism. Hormone replacement therapy (HRT) could be a beneficial in female patients with JSLE to improve their lipid profile by increasing HDL. Previously, this view has been controversial since exogenous estrogen could induce disease symptoms and cause flares. Evidence surrounding this however remains elusive and further investigation is required [59]. . Further hormone studies will be required to underpin the exact signalling mechanisms that become defective in JSLE patients.

Relative to effector T-cell subsets, Tregs rely more heavily on lipids for their function via beta-oxidation [49]. Treg frequency is increased in individuals with dyslipidaemia; a positive correlation between Treg frequency and ApoB, LDL cholesterol and total cholesterol was identified [50]. In contrast, Treg frequency was increased in HDL-treated mice and a positive correlation between Treg frequency and HDL-cholesterol was reported in statin treated adults [51]. This study also showed that ABCG1 expression (implicated in cholesterol efflux) in human peripheral blood Tregs correlated with an increased Treg frequency; it was concluded that accumulation of cholesterol could alter T-cell homeostasis. Another study identified that depletion of Tregs in atherosclerosis-prone, low-density lipoprotein receptor-deficient (Ldlr-/-) mice resulted in increased atherosclerosis associated with elevated plasma cholesterol and VLDL due to lack of clearance [52]. This demonstrates that Tregs may elicit the capability to alter lipoprotein metabolism. Differences in the Treg:lipoprotein axis in males and females could contribute to the differential Treg frequency and function observed here and could play a significant role in the development of atherosclerosis in females in the context of autoimmunity.

This study represents a rare opportunity to investigate Treg phenotype and serum metabolome in a young age matched JSLE cohort. Most studies have found a reduction in Treg frequency in SLE, with an improvement after treatment [56], thus it was important that there was no significant difference in treatment between males and females in the cohort. Importantly we found no difference in the lipid metabolome of patients in this cohort with regards to steroid treatment despite evidence that steroid use can alter plasma lipid levels [57, 58]. Sex differences in Treg phenotype and function and serum metabolomics were lost in JSLE. Together this work adds complexity to the study of JSLE pathogenesis but highlights the importance of considering sex in autoimmune and metabolic research.

In summary, here we have identified a novel link between lipoprotein metabolism and Treg function in males and females. Males have a more pro-atherogenic metabolome and this influences Tregs to display a more suppresive phenotype compared to females; this could help to explain differences in autoimmune susceptibility. In JSLE, these sex-associated differences in Treg phenotype and serum metabolic profile are lost. This may be associated with altered hormone signalling although further investigation is required to dissect the precise mechanisms.

## MATERIALS AND METHODS

### Patients and control samples

Peripheral blood was collected from 35 JSLE patients (12 male, median age 18, 22 female, median age 20, all puberty tanner stage 4-5) attending a young adult or adolescent rheumatology clinic at University College London Hospital (UCLH) or Great Ormond Street Hospital (GOSH) respectively (Supplementary Table 1). Blood from 39 age, ethnicity and sex matched teenage and young adult healthy controls (HC) was also collected (17 males, median age 18, 22 females, median age 18, all puberty tanner stage 4-5). 20 Gender dysphoria blood samples were collected from individuals (XY, n=10, median age 18 and XX, n=10, median age 18) attending the UCLH gender dysphoria endocrine clinic while their routine bloods were taken for their treatment. Informed written consent was acquired from patients and healthy donors under the ethical approval reference: REC11/LO/0330. Questionnaires provided the tanner puberty stage of all donors as well as their current use of contraception. All information was stored as anonymised data. Detailed clinical characteristics and disease features were recorded from patient files and questionnaires. Disease activity was calculated using the SLE Disease Activity Index (SLEDAI). Donor anonymous 50ml leukocyte blood cones were obtained from NHS Blood and Transplant (NHSBT) (Colindale) (ethical approval reference: REC 16/YH/0306).

### Flow cytometry

#### Surface Staining

10^6^ PBMCs were stained with fixable blue dead cell stain (ThermoFisher) or Zombie NIR™ Fixable Viability Kit (Biolegend) in PBS followed by washes and surface marker staining in Brilliant™ Stain Buffer (BD). The following antibodies were used: CD3-BV786 (Biolegend), CD8-BV421 (Biolegend), CD4-BUV395 (BD), CD25-PeDazzle594 (Biolegend), CD127-BV711 (Biolegend), CD27-AF700 (Biolegend) and CD27-PECy7 (Biolegend) (Supplementary Figure 1 for gating). 1 million PBMCs per sample were stained for 30mins on ice followed by subsequent washes and fixation in 2% PFA. In some cases cells were also surface stained with GLUT-1-AF647 (abcam), CD69-BV510 (Biolegend) and PD-1-APC-Cy7 (Biolegend) antibodies using the same method.

#### Lipid staining

In some cases following surface staining, cells were subsequently stained with, cholera toxin B-FITC (CTB) binds to the glycosphingolipid GM1 [62], (Sigma, 1:100 in PBS) for 30 mins on ice. Cells were fixed with 2% PFA (30mins at room temperature) followed by membrane cholesterol staining with filipin complex from (Sigma, 50μg/ml in PBS) [62] for 2hrs at room temperature.

For membrane order analysis, live cells were stained with 4 uM di-4-ANEPPDHQ (ANE) (ThermoFisher Scientific) for flow cytometry analysis as previously described [25].

#### Intracellular staining

PBMCs were cultured for 4.5 hours in PMA (50ng/ml), ionomycin (250ng/ml) (Sigma) and GolgiPlug™ Protein Transport Inhibitor (BD Biosciences) followed by staining with Zombie NIR™ Fixable Viability Kit (Biolegend) (1/1000 in PBS). Surface staining was carried out as above. Cells were then fixed and permeabilised in Fix/Perm buffer (Biolegend) (overnight at 4°C) followed by intracellular cytokine staining with anti-human IL-10-APC (BD Biosciences), IL-4-PERCP Cy5.5 (Biolegend) and IFNγ-eFlour450 (ebioscience) in permeabilisation wash buffer (Biolegend) (45mins on ice). Staining for FOXP3-PE (Biolegend), T-bet-APC (Biolegend) and GATA-3-AP488 (Biolegend) was carried out using the same method but without stimulation.

#### Data acquisition and analysis

Data was acquired using a BD LSRFORTESSA X-20 flow cytometer (1×10^6^-3×10^6^ cells/sample) and analysis was performed using FlowJo Single Cell Analysis Software (TreeStar). Application settings were created to begin with and Cytometer Setup and Tracking (CS&T) (BD) beads were run on the flow cytometer before each session to assess cytometer performance. Application settings were applied to panel templates each time prior to compensation to ensure that all immunophenotyping data was comparable over time. Gating strategies are shown in Supplementary Figure 1. Absolute cell counts were calculated by multiplying the percentage of the population of interest in the total lymphocyte count by the average UCLH reference guidelines for healthy donor lymphocyte count (1.225×10^9^ cells/litre).

#### Cell sorting

PBMCs were washed in MACS buffer (PBS (Sigma), 2% FBS (Labtech) and 1mM EDTA (Sigma)) and stained for 30 minutes with anti-human CD4-BUV395 (BD Biosciences), CD25-PE-Dazzle594 (Biolegend), CD127-BV711 (Biolegend) and CD14-APC (Biolegend) and sorted using a BD FACSAria cell sorter into collection media (1xPBS, 20% FBS). Purity checks were completed on sorted cells (Supplementary Figure 2).

### Cell Culture

#### Suppression Assay

FACS sorted Tregs, Tresp and monocytes were used for suppression assays. Treg purity was confirmed by FOXP3 intracellular staining (Supplementary Figure 2). Tresps (CD25^−^CD127^+^) were labelled for 20 minutes at room temperature with Cell Trace Violet (CTV, thermofisher) in PBS and cultured at a density of 8×10^4^ cells/well in RPMI 1640 medium (Thermofisher) supplemented with 10% FBS (ThermoFisher), Penicillin (100 IU/ml) and Streptomycin (100 μg/ml) (Gibco) with 1ug/mL soluble anti-CD3 (OKT3 mAb) (ebioscience) and 1ug/ml soluble anti-CD28 (ebioscience) in a 96 well flat bottom plate. Cells were co-cultured with unlabelled Treg cells at equal and descending density as well as monocytes at set density of 8×10^4^. Proliferating cells expressing CTV were fixed in PFA and measured by flow cytometry after 4 days. Additional surface staining of CD4 was carried out to ensure monocytes were excluded from the analysis (Supplemental Figure 3).

#### LDL/VLDL isolation from serum and cell culture

LDL/VLDL was isolated from 2-3ml pooled healthy male or female serum using LDL/VLDL Purification Kit (Ultracentrifugation Free) (CELL BIOLABS) according to manufacturer’s instructions. Purified LDL/VLDL was dialysed in 10ml PBS in Amicon Ultra centrifugal filter units (MILLIPORE). A Pierce™ Bicinchoninic Acid assay (BCA) Protein Assay Kit (ThermoFisher) was used to quantify the isolated LDL/VLDL (Apolipoprotein B protein expression). Isolated lipoproteins were filtered through a 0.22um filter and stored at 4 °C for a maximum of 6 weeks. 1×10^6^ PBMCs were cultured with isolated VLDL (50ug/ul) in RPMI-1640 medium or an equal volume of lipoprotein depleted FBS (Kalen Biomedical) for 24 hours.

#### Phospholipid detection

For phospholipid fluorescent staining, the HCS LipidTOX™ Phospholipidosis and Steatosis Detection Kit (thermofisher) was used as by the manufacturer’s instruction. Here the phospholipid stain was added to the culture with the VLDL stimulus and cells were cultured for 24 hours.

### RNA extraction and analysis

*Ex-vivo* FACS sorted cell subsets or *in vitro* cultured CD4^+^ T-cell subsets at density of 0.5-1×10^6^ were lysed and RNA was extracted and isolated using the PicoPure RNA Isolation Kit (ThermoFisher) according to manufacturer’s instructions. cDNA was synthesised using qScript cDNA SuperMix (Quantabio). PerfeCTa^®^ SYBR^®^ Green FastMix (VWR) was added to cDNA for fluorescence quantification using a Stratagene Mx3000P qPCR System (Agilent Technologies). Forward and reverse primers were designed using ncbi PrimerBlast [63] and ensemble (https://www.ensembl.org/index.html) and were used at 100nM in each reaction: Cyclophilin A (forward, GCATACGGGTCCTGGCATCTTGTCC; reverse, ATGGTGATCTTCTTGCTGGTCTTGC); UDP-Glucose Ceramide Glucosyltransferase (UGCG) (forward, CGTCCTCTTCTTGGTGCTGT; reverse, AGAGAGACACCTGGGAGCTT); ATP-binding cassette transporter A1 (ABCA1) (forward, TGAGCTACCCACCCTATGAACA; reverse, CCCCTGAACCCAAGGAAGTG); ATP-binding cassette transporter G1 (ABCG1) (forward, TGCAATCTTGTGCCATATTTGA; reverse, CCAGCCGACTGTTCTGATCA); 3-hydroxy-3-methyl-glutaryl-coenzyme A reductase (HMGCR) (forward, CAGCCATTTTGCCCGAGTTT; reverse, AGCGACTGTGAGCATGAACA). Data was collected by MxPro software (Agilent Technologies) and analysis was carried out using Microsoft Excel 2010 where all data was normalised to reference gene cyclophilin A. The Delta- or Delta-Delta-Ct (dCt or ddCt) algorithmic method of quantification was used to determine relative gene expression [64].

### Metabolomics

Nightingale metabolomics (https://nightingalehealth.com/), a service that measures 220 blood biomarkers [65], was used to measure biomarkers in healthy control and JSLE patient serum (Supplemental Table 2 for list of biomarkers).

### Statistical analysis

Analysis was performed in GraphPad Prism 7 software. Data was tested for normal distribution using Kolmogorov-Smirnov test and parametric or non-parametric tests were used accordingly. Unpaired or pared t-tests were used to compare ex-vivo patient group data or culture/pre-post matched sample data respectively. Linear regression was performed with a 95% confidence interval used to calculate significance (Pearson correlation). Multiple testing was also accounted for by use of FDR correction (Benjamini, Krieger and Yekutieli approach) of p-value thresholds. This was carried out on phenotyping and metabolomics data using Prism 7 software analysis of p-value stacks. MultiExperiment Viewer (MeV) was used to produce heat maps and perform hierarchical clustering (Pearson correlation) on large data sets. Clustering was performed on z-score transformed raw data values (*the signed number of standard deviations by which the value of an observation or data point differs from the mean value of what is being observed or measured*). These were calculated in Microsoft Excel 2010 using the equation: *((Individual Value - Population Mean) / (Population Standard Deviation – Square Root of the Total Sample Number))*.

Clustvis (https://biit.cs.ut.ee/clustvis/) web tool was used to carry out principle component analysis (PCA) of metabolomics data.

## Contributors

Design of research study; ECJ, IPT, CC, YI. Acquiring data; GAR, KW, MA, AR, HP, CW. Recruiting patients; HP, YI, CC, CW, DAI. Analyzing data; GAR Writing the manuscript; GAR, ECJ: Review of the manuscript; DAI, CC, IPT. All authors approved the final version.

## Acknowledgments

GAR was supported by a PhD studentship from Lupus UK and The Rosetrees Trust (M409). KEW was funded by a British Heart Foundation PhD Studentship (FS/13/59/30649). This work was supported by the Adolescent Centre at UCL UCLH and GOS funded by Arthritis Research UK (20164 and 21953), Great Ormond Street Children’s Charity and the NIHR Biomedical Research Centres at GOSH and UCLH. The views expressed are those of the authors and not necessarily those of the NHS, the NIHR or the Department of Health.

## References

1. Klein, S.L. and K.L. Flanagan, Sex differences in immune responses. Nat Rev Immunol, 2016. 16(10): p. 626–638.

2. Uppal, S.S., S. Verma, and P.S. Dhot, Normal values of CD4 and CD8 lymphocyte subsets in healthy Indian adults and the effects of sex, age, ethnicity, and smoking. Cytometry Part B-Clinical Cytometry, 2003. 52B(1): p. 32–36.

3. Sankaran-Walters, S., et al., Sex differences matter in the gut: effect on mucosal immune activation and inflammation. Biology of Sex Differences, 2013. 4: p. 12.

4. Lisse, I.M., et al., T-lymphocyte subsets in West African children: Impact of age, sex, and season. Journal of Pediatrics, 1997. 130(1): p. 77–85.

5. Hughes, G.C. and D. Choubey, Modulation of autoimmune rheumatic diseases by oestrogen and progesterone. Nat Rev Rheumatol, 2014. 10(12): p. 740–751.

6. Kone-Paut, I., et al., Lupus in adolescence. Lupus, 2007. 16(8): p. 606–12.

7. Ambrose, N., et al., Differences in disease phenotype and severity in SLE across age groups. Lupus, 2016.

8. Urowitz, M.B., et al., The bimodal mortality pattern of systemic lupus erythematosus. Am J Med, 1976. 60(2): p. 221–5.

9. Michel, V. and M. Bakovic, Lipid rafts in health and disease. Biol Cell, 2007. 99(3): p. 129–40.

10. Matthews, K.A., et al., MENOPAUSE AND RISK-FACTORS FOR CORONARY HEART-DISEASE. New England Journal of Medicine, 1989. 321(10): p. 641–646.

11. Abbey, M., et al., Effects of menopause and hormone replacement therapy on plasma lipids, lipoproteins and LDL-receptor activity. Maturitas, 1999. 33(3): p. 259–269.

12. Kannel, W.B., CITATION CLASSIC - SERUM-CHOLESTEROL, LIPOPROTEINS, AND THE RISK OF CORONARY HEART-DISEASE - THE FRAMINGHAM-STUDY. Current Contents/Life Sciences, 1983(29): p. 18–18.

13. Xing, Q., et al., Elevated Th17 cells are accompanied by FoxP3+Treg cells decrease in patients with lupus nephritis. Rheumatology International, 2012. 32(4): p. 949–958.

14. Ma, J.L., et al., The imbalance between regulatory and IL-17-secreting CD4(+) T cells in lupus patients. Clinical Rheumatology, 2010. 29(11): p. 1251–1258.

15. Yang, J., et al., Th17 and Natural Treg Cell Population Dynamics in Systemic Lupus Erythematosus. Arthritis and Rheumatism, 2009. 60(5): p. 1472–1483.

16. Afshan, G., N. Afzal, and S. Qureshi, CD4(+)CD25(hi) Regulatory T Cells in Healthy Males and Females Mediate Gender Difference in the Prevalence of Autoimmune Diseases. Clinical Laboratory, 2012. 58(5-6): p. 567–571.

17. Collier, F.M., et al., The ontogeny of naive and regulatory CD4(+) T-cell subsets during the first postnatal year: a cohort study. Clinical & Translational Immunology, 2015. 4: p. 8.

18. Tai, P., et al., Induction of regulatory T cells by physiological level estrogen. Journal of Cellular Physiology, 2008. 214(2): p. 456–464.

19. Arruvito, L., et al., Expansion of CD4(+)CD25(+) and FOXP3(+) regulatory T cells during the follicular phase of the menstrual cycle: Implications for human reproduction. Journal of Immunology, 2007. 178(4): p. 2572–2578.

20. Tommasini, A., et al., X-chromosome inactivation analysis in a female carrier of FOXP3 mutation. Clinical and Experimental Immunology, 2002. 130(1): p. 127–130.

21. Klein, S.L. and K.L. Flanagan, Sex differences in immune responses. Nature Reviews Immunology, 2016. 16: p. 626.

22. Zhu, J.F., et al., GATA-3 promotes Th2 responses through three different mechanisms: induction of Th2 cytokine production, selective growth of Th2 cells and inhibition of Th1 cell-specific factors. Cell Research, 2006. 16(1): p. 3–10.

23. McDonald, G., et al., Normalizing glycosphingolipids restores function in CD4(+) T cells from lupus patients. Journal of Clinical Investigation, 2014. 124(2): p. 712–724.

24. Waddington, K.E., et al., Activation of the liver X receptors alters CD4^+^ T cell membrane lipids and signalling through direct regulation of glycosphingolipid synthesis. bioRxiv, 2019: p. 721050.

25. Miguel, L., et al., Primary Human CD4(+) T Cells Have Diverse Levels of Membrane Lipid Order That Correlate with Their Function. Journal of Immunology, 2011. 186(6): p. 3505–3516.

26. Allende, M.L. and R.L. Proia, Lubricating cell signaling pathways with gangliosides. Current Opinion in Structural Biology, 2002. 12(5): p. 587–592.

27. Freedman, D.S., et al., Sex and Age Differences in Lipoprotein Subclasses Measured by Nuclear Magnetic Resonance Spectroscopy: The Framingham Study. Clinical Chemistry, 2004. 50(7): p. 1189–1200.

28. Freedman, D.S., et al., Distribution and correlates of high-density lipoprotein subclasses among children and adolescents. Metabolism-Clinical and Experimental, 2001. 50(3): p. 370–376.

29. Fonseca, M.J., et al., Association of Pubertal Development With Adiposity and Cardiometabolic Health in Girls and Boys-Findings From the Generation XXI Birth Cohort. The Journal of adolescent health : official publication of the Society for Adolescent Medicine, 2019.

30. Mascarenhas, L.P.G., et al., Variability of lipid and lipoprotein concentrations during puberty in Brazilian boys. Journal of Pediatric Endocrinology & Metabolism, 2015. 28(1-2): p. 125–131.

31. Eissa, M.A., et al., Changes in Fasting Lipids during Puberty. Journal of Pediatrics, 2016. 170: p. 199–205.

32. Storey, S.M., et al., Selective cholesterol dynamics between lipoproteins and caveolae/lipid rafts. Biochemistry, 2007. 46(48): p. 13891–13906.

33. Miyara, M., et al., Functional Delineation and Differentiation Dynamics of Human CD4(+) T Cells Expressing the FoxP3 Transcription Factor. Immunity, 2009. 30(6): p. 899–911.

34. Dr Michael W Beresford, L.U. Juvenile-Onset Lupus. [cited 2016; Available from: http://www.lupusuk.org.uk/medical/gp-guide/diagnosis-of-lupus/associated-illnesses/childhood/.

35. Brunkow, M.E., et al., Disruption of a new forkhead/winged-helix protein, scurfin, results in the fatal lymphoproliferative disorder of the scurfy mouse. Nature Genetics, 2001. 27(1): p. 68–73.

36. Halim, L., et al., An Atlas of Human Regulatory T Helper-like Cells Reveals Features of Th2-like Tregs that Support a Tumorigenic Environment. Cell Reports, 2017. 20(3): p. 757–770.

37. Wang, Y.Q., M.A. Su, and Y.S.Y. Wan, An Essential Role of the Transcription Factor GATA-3 for the Function of Regulatory T Cells. Immunity, 2011. 35(3): p. 337–348.

38. Wohlfert, E.A., et al., GATA3 controls Foxp3(+) regulatory T cell fate during inflammation in mice. Journal of Clinical Investigation, 2011. 121(11): p. 4503–4515.

39. Nazarov-Stoica, C., et al., CD28 Signaling in T Regulatory Precursors Requires p56(Ick) and Rafts Integrity to Stabilize the Foxp3 Message. Journal of Immunology, 2009. 182(1): p. 102–110.

40. Balamuth, F., et al., Distinct patterns of membrane microdomain partitioning in Th1 and Th2 cells. Immunity, 2001. 15(5): p. 729–738.

41. Surls, J., et al., Increased Membrane Cholesterol in Lymphocytes Diverts T-Cells toward an Inflammatory Response. 2012, Figshare.

42. Cook, K.D. and J. Miller, TCR-Dependent Translational Control of GATA-3 Enhances Th2 Differentiation. Journal of Immunology, 2010. 185(6): p. 3209–3216.

43. Watson, L., et al., Disease activity, severity, and damage in the UK Juvenile-Onset Systemic Lupus Erythematosus Cohort. Arthritis Rheum, 2012. 64(7): p. 2356–65.

44. Lohner, S., et al., Gender Differences in the Long-Chain Polyunsaturated Fatty Acid Status: Systematic Review of 51 Publications. Annals of Nutrition and Metabolism, 2013. 62(2): p. 98–112.

45. Wang, Q., et al., Effects of hormonal contraception on systemic metabolism: cross-sectional and longitudinal evidence. International Journal of Epidemiology, 2016. 45(5): p. 1445–1457.

46. Fisher, A.D., Cross-sex hormone therapy in trans persons is safe and effective at short-time follow-up: results from the European Network for the Investigation of Gender Incongruence (vol 11, pg 1999, 2014). Journal of Sexual Medicine, 2016. 13(4): p. 732–732.

47. Jarin, J., et al., Cross-Sex Hormones and Metabolic Parameters in Adolescents With Gender Dysphoria. Pediatrics, 2017. 139(5).

48. Della Torre, S., et al., An Essential Role for Liver ERα in Coupling Hepatic Metabolism to the Reproductive Cycle. Cell Reports. 15(2): p. 360–371.

49. Gerriets, V.A. and J.C. Rathmell, Metabolic pathways in T cell fate and function. Trends in Immunology, 2012. 33(4): p. 168–173.

50. Guasti, L., et al., Relationship between regulatory T cells subsets and lipid profile in dyslipidemic patients: a longitudinal study during atorvastatin treatment. Bmc Cardiovascular Disorders, 2016. 16: p. 9.

51. Rueda, C.M., et al., High density lipoproteins selectively promote the survival of human regulatory T cells. Journal of Lipid Research, 2017. 58(8): p. 1514–1523.

52. Klingenberg, R., et al., Depletion of FOXP3(+) regulatory T cells promotes hypercholesterolemia and atherosclerosis. Journal of Clinical Investigation, 2013. 123(3): p. 1323–1334.

53. Sampedro, M.C., et al., VLDL modulates the cytokine secretion profile to a proinflammatory pattern. Biochemical and Biophysical Research Communications, 2001. 285(2): p. 393–399.

54. Tan, T.C., et al., Differences between Male and Female Systemic Lupus Erythematosus in a Multiethnic Population. Journal of Rheumatology, 2012. 39(4): p. 759–769.

55. Crosslin, K.L. and K.L. Wiginton, Sex Differences in Disease Severity Among Patients With Systemic Lupus Erythematosus. Gender Medicine, 2011. 8(6): p. 365–371.

56. Horwitz, D.A., Regulatory T cells in systemic lupus erythematosus: past, present and future. Arthritis Research & Therapy, 2008. 10(6): p. 9.

57. Ettinger, W.H. and W.R. Hazzard, PREDNISONE INCREASES VERY LOW-DENSITY LIPOPROTEIN AND HIGH-DENSITY LIPOPROTEIN IN HEALTHY-MEN. Metabolism-Clinical and Experimental, 1988. 37(11): p. 1055–1058.

58. Elshaboury, A.H. and T.M. Hayes, HYPERLIPIDEMIA IN ASTHMATIC-PATIENTS RECEIVING LONG-TERM STEROID-THERAPY. Bmj-British Medical Journal, 1973. 2(5858): p. 85–86.

59. Holroyd, C.R. and C.J. Edwards, The effects of hormone replacement therapy on autoimmune disease: rheumatoid arthritis and systemic lupus erythematosus. Climacteric, 2009. 12(5): p. 378–386.

60. Maselli, A., et al., Low expression of estrogen receptor β in T lymphocytes and high serum levels of anti-estrogen receptor α antibodies impact disease activity in female patients with systemic lupus erythematosus. Biology of Sex Differences, 2016. 7: p. 3.

61. Page, S.T., et al., Effect of medical castration on CD4(+) CD25(+) T cells, CD8(+) T cell IFN-gamma expression, and NK cells: a physiological role for testosterone and/or its metabolites. American Journal of Physiology-Endocrinology and Metabolism, 2006. 290(5): p. E856–E863.

62. Waddington, K.E., I. Pineda-Torra, and E.C. Jury, Analyzing T-Cell Plasma Membrane Lipids by Flow Cytometry. Methods in molecular biology (Clifton, N.J.), 2019. 1951: p. 209–216.

63. primer-blast, n. http://www.ncbi.nlm.nih.gov/tools/primer-blast/. [cited 2015/16.

64. Livak, K.J. and T.D. Schmittgen, Analysis of relative gene expression data using real-time quantitative PCR and the 2(T)(-Delta Delta C) method. Methods, 2001. 25(4): p. 402–408.

65. Robinson, G.A., et al., Metabolomics in juvenile-onset SLE: identifying new biomarkers to predict cardiovascular risk. medRxiv, 2019: p. 19000356.

