## Supplementary material for "Sex differences in autoimmunity could be associated with altered regulatory T cell phenotype and lipoprotein metabolism"

### Supplemental Tables

Supplemental Table 1: Demographic comparison between adolescent healthy post-pubertal males and females within the cohort

| Demographic | Male | Female | P value* |
|---|---|---|---|
| **Number** | 17 | 22 | - |
| **Median Age (range)** | 18 (16-25) | 17.5 (16-25) | $^{+}$0.85 |
| **Ethnicity** | | | |
| **White** | 8 (47%) | 12 (55%) | *0.75 |
| **Asian** | 6 (35%) | 3 (18%) | *0.14 |
| **Black** | 1 (6%) | 2 (9%) | *1.00 |
| **Other** | 2 (12%) | 4 (18%) | *0.68 |

*Fisher's exact test or $^{+}$unpaired t-test was used

#### Supplemental Table 2: List of serum metabolic biomarkers

| | | |
|---|---|---|
| • Cholesterol | • Lipoprotein subclasses | • M-LDL-CE |
| • Total-C | • XXL-VLDL-P | • M-LDL-FC |
| • VLDL-C | • XXL-VLDL-L | • M-LDL-TG |
| • Remnant-C | • XXL-VLDL-PL | • S-LDL-P |
| • LDL-C | • XXL-VLDL-C | • S-LDL-L |
| • HDL-C | • XXL-VLDL-CE | • S-LDL-PL |
| • HDL2-C | • XXL-VLDL-FC | • S-LDL-C |
| • HDL3-C | • XXL-VLDL-TG | • S-LDL-CE |
| • Esterified-C | • XL-VLDL-P | • S-LDL-FC |
| • Free-C | • XL-VLDL-L | • S-LDL-TG |
| • Glycerides and phospholipids | • XL-VLDL-PL | • XL-HDL-P |
| • Total triglycerides | • XL-VLDL-C | • XL-HDL-L |
| • VLDL-TG | • XL-VLDL-CE | • XL-HDL-PL |
| • LDL-TG | • XL-VLDL-FC | • XL-HDL-C |
| • HDL-TG | • XL-VLDL-TG | • XL-HDL-CE |
| • Phosphoglycerides | • L-VLDL-P | • XL-HDL-FC |
| • TG/PG | • L-VLDL-L | • XL-HDL-TG |
| • Total cholines | • L-VLDL-PL | • L-HDL-P |
| • Phosphatidylcholines | • L-VLDL-C | • L-HDL-L |
| • Sphingomyelins | • L-VLDL-CE | • L-HDL-PL |
| • Apolipoproteins | • L-VLDL-FC | • L-HDL-C |
| • ApoB | • L-VLDL-TG | • L-HDL-CE |
| • ApoA1 | • M-VLDL-P | • L-HDL-FC |
| • ApoB/ApoA1 | • M-VLDL-L | • L-HDL-TG |
| • Fatty acids | • M-VLDL-PL | • M-HDL-P |
| • FAw3/FA | • M-VLDL-C | • M-HDL-L |
| • FAw6/FA | • M-VLDL-CE | • M-HDL-PL |
| • PUFA/FA | • M-VLDL-FC | • M-HDL-C |
| • MUFA/FA | • M-VLDL-TG | • M-HDL-CE |
| • SFA/FA | • S-VLDL-P | • M-HDL-FC |
| • DHA/FA | • S-VLDL-L | • M-HDL-TG |
| • LA/FA | • S-VLDL-PL | • S-HDL-P |
| • Amino acids | • S-VLDL-C | • S-HDL-L |
| • Alanine | • S-VLDL-CE | • S-HDL-PL |
| • Glutamine | • S-VLDL-FC | • S-HDL-C |
| • Glycine | • S-VLDL-TG | • S-HDL-CE |
| • Histidine | • XS-VLDL-P | • S-HDL-FC |
| • Isoleucine | • XS-VLDL-L | • S-HDL-TG |
| • Leucine | • XS-VLDL-PL | |
| • Valine | • XS-VLDL-C | |
| • Phenylalanine | • XS-VLDL-CE | |
| • Tyrosine | • XS-VLDL-FC | |
| • Glycolysis related metabolites | • XS-VLDL-TG | |
| • Glucose | • IDL-P | |
| • Lactate | • IDL-L | |
| • Pyruvate | • IDL-PL | |
| • Citrate | • IDL-C | |
| • Glycerol | • IDL-CE | |
| • Ketone bodies | • IDL-FC | |
| • Acetate | • IDL-TG | |
| • Acetoacetate | • L-LDL-P | |
| • 3-hydroxybutyrate | • L-LDL-L | |
| • Fluid balance | • L-LDL-PL | |
| • Creatinine | • L-LDL-C | |
| • Albumin | • L-LDL-CE | |
| • Inflammation | • L-LDL-FC | |
| • Glycoprotein acetyls | • L-LDL-TG | |
| • Lipoprotein particle sizes | • M-LDL-P | |
| • VLDL particle size | • M-LDL-L | |
| • LDL particle size | • M-LDL-PL | |
| • HDL particle size | • M-LDL-C | |

**Supplemental Table 2: List of serum metabolic biomarkers.** Nightingale Health metabolomics platform (https://nightingalehealth.com/) was used to measure biomarkers in HC and JSLE patient serum. This service measures blood metabolic biomarkers using nuclear magnetic resonance (NMR) spectroscopy. The platform can simultaneously measure amino acids, fatty acids, glycolysis metabolites, routine lipid measures (mmol/l), apolipoproteins (g/l) and in depth lipoprotein measurements such as particle size (nm), concentration and lipid content (mmol/l). The platform provides repeatable measurements with no batch effects. The Nightingale Health service has been thoroughly validated and has been used to measure over 500,000 samples from both research and clinical trials.

Abbreviations: Apo, apolipoprotein; VLDL, very low density lipoprotein; IDL, intermediate density lipoprotein; LDL, low density lipoprotein; HDL, high density lipoprotein; XXL-VLDL, chylomicrons and extremely large VLDL; X-Large, very large; X-small, very small; Est (esterified), PG (Phosphoglyceride), PC, Phosphatidylcholine; SM, Sphingomyelins; Unsat, Unsaturated; DHA, Docosahexaenoic acid; LA, Linoleic acid; FAw3, Omega-3 fatty acids; FAw6, Omega-6 fatty acids; PUFA, Polyunsaturated fatty acids; MUFA, Monounsaturated fatty acids; SFA, Saturated fatty acids; TG, triglycerides; PL, phospohlipids, FC, free cholesterol, C, cholesterol, CE, cholesterol esters; P, particle; L, lipid.

#### Supplemental Table 3: Metabolites significantly associated with adolescent healthy post-pubertal males and females

| Metabolite | Male, Mean [mmol/L] (range) | Female, Mean[mmol/L] (range) | P value |
|---|---|---|---|
| **Male associated:** | | | |
| XXL-VLDL-Particles | 1.10E-10 (0 - 3.64E-10) | 2.90E-11 (0 - 1.80E-10) | 0.0106 |
| XXL-VLDL- total lipids | 2.33E-2 (0 - 7.75E-02) | 6.08E-3 (0 - 3.83E-2) | 0.0109 |
| XXL-VLDL- phospholipids | 2.59E-03 (0 - 9.27E-03) | 5.72E - 04 (0-4.38E-03) | 0.0115 |
| XXL-VLDL- cholesterol | 3.53E-03 (0 - 1.28E-02) | 9.21E - 04 (0-6.29E-03) | 0.0166 |
| XXL-VLDL- cholesterol ester | 1.88E-03 (0 - 6.78E-03) | 5.62E-04 (0 - 3.57E-03) | 0.0233 |
| XXL-VLDL- free cholesterol | 1.65E-03 (0 - 5.98E-03) | 3.60E-04 (0 - 2.73E-03) | 0.0115 |
| XXL-VLDL- triglycerides | 1.71E-02 (0 - 5.55E-02) | 4.59E-03 (0 - 2.76E-02) | 0.0099 |
| XL-VLDL- Particles | 6.312-10 (0 - 2.07E-09) | 1.51E-10 (0 - 9.99e-10) | 0.0072 |
| XL-VLDL- total lipids | 6.07E-02 (0 - 2.00E-01) | 1.45E-02 (0 - 9.65E-02) | 0.0075 |
| XL-VLDL- phospholipids | 9.10E-03 (0 - 3.23E-02) | 2.04E-03 (0 - 1.49E-02) | 0.0104 |
| XL-VLDL- cholesterol | 1.09E-02 (0 - 3.81E-02) | 2.66E-03 (0 - 1.85E-02) | 0.0109 |
| XL-VLDL- cholesterol ester | 6.33E-03 (0 - 2.08E-02) | 1.67E-03 (0 - 1.08E-02) | 0.0093 |
| XL-VLDL- free cholesterol | 4.58 (0 - 1.73E-02) | 9.89E-04 (0 - 7.67E-03) | 0.0143 |
| XL-VLDL- triglycerides | 4.06E-02 (0 - 1.30E-01) | 9.78E-03 (0 - 6.32E-02) | 0.0064 |
| L-VLDL- Particles | 4.39E-09 (0 - 1.24E-08) | 1.46E-09 (0 - 6.26E-09) | 0.0058 |
| L-VLDL- total lipids | 2.51E-01 (0 - 7.14E-01) | 8.26E-02 (0 - 3.61E-01) | 0.0059 |
| L-VLDL- phospholipids | 4.40E-02 (0 - 1.27E-01) | 1.44E-02 (0 - 6.36E-02) | 0.0065 |
| L-VLDL- cholesterol | 5.32E-02 (0 - 1.56E-01) | 1.70E-02 (0 - 8.17E-02) | 0.007 |
| L-VLDL- cholesterol ester | 2.92E-02 (0 - 7.94E-02) | 1.09E-02 (0 - 4.56E-02) | 0.0088 |
| L-VLDL- free cholesterol | 2.40E-02 (0 - 7.61E-02) | 6.13E-03 (0 - 3.61E-02) | 0.0062 |
| L-VLDL- triglycerides | 1.54E-01 (0 - 4.32E-01) | 5.13E-02 (0 - 2.16E-01) | 0.0054 |
| M-VLDL- Particles | 1.60E-08 (4.60E-09 - 3.49E-08) | 8.36E-09 (0 - 2.02E-08) | 0.0047 |
| M-VLDL- total lipids | 5.27E-01 (1.49E-01 - 1.15) | 2.77E-01 (0 - 2.02E-08) | 0.0053 |
| M-VLDL- phospholipids | 1.04E-01 (3.29E-02 - 2.21E-01) | 5.71E-02 (0 - 1.32E-01) | 0.0063 |
| M-VLDL- cholesterol | 1.24E-01 (2.24E-02 - 2.76E-01) | 7.18E-02 (0 - 1.72E-01) | 0.0225 |
| M-VLDL- free cholesterol | 5.69E-02 (9.84E-03 - 1.34E-01) | 2.72E-02 (0 - 7.52E-02) | 0.0071 |
| M-VLDL- triglycerides | 2.99E-01 (8.22E-02 - 6.55E-01) | 1.48E-01 (0 - 3.65E-01) | 0.0025 |
| S-VLDL- Particles | 2.40E-08 (1.17E-08 - 4.00E-08) | 1.71E-08 (7.91E-09 - 2.87E-08) | 0.017 |
| S-VLDL- total lipids | 4.58E-01 (2.24E-01 - 7.56E-01) | 3.32E-01 (1.54E-01 - 5.52E-01) | 0.0231 |
| S-VLDL- phospholipids | 1.08E-01 (5.88E-02 - 1.65E-01) | 8.17E-02 (4.56E-02 - 1.30E-01) | 0.0203 |
| S-VLDL- free cholesterol | 5.90E-02 (2.78E-02 - 9.40E-02) | 4.42E-02 (1.96E-02 - 7.20E-02) | 0.0396 |
| S-VLDL- triglycerides | 2.16E-01 (9.44E-02 - 3.79E-01) | 1.37E-01 (5.13E-02 - 2.52E-01) | 0.0038 |
| XS-VLDL- triglycerides | 8.44E-02 (4.25E-02 - 1.23E-01) | 6.61E-02 (3.44E-02 - 1.04E-01) | 0.0326 |
| VLDL-Diameter | 3.76E+01 (3.49E+01 - 4.00E+1) | 3.58E+01 (3.37E+01 - 3.81E+01) | 0.0003 |
| VLDL- triglycerides | 8.16E-01 (2.27E-01 - 1.77) | 4.26E-01 (1.05E-01 - 1.03) | 0.0036 |
| Serum-Triglycerides | 1.12 (4.30E-01 - 2.15) | 7.19E-01 (3.07E-01 - 1.42) | 0.0092 |
| Monounsaturated fatty acids | 2.70E+01 (2.15E+01 - 3.12E+01) | 2.49E+01 (2.14E+01 - 2.84E +01) | 0.0418 |
| Saturated fatty acids | 3.55E+01 (3.14E+01 - 3.89E+01) | 3.43E+01 (3.14E+01 - 3.65E+01) | 0.0458 |
| Isoleucine | 5.89E-02 (3.65E-02 - 1.08E-01) | 4.35E-02 (2.50E-02 - 8.34E-02) | 0.0157 |
| Leucine | 7.76E-02 (5.77E-02 - 1.25E-01) | 7.76E-02 (3.90E-02 - 9.86E-02) | 0.0057 |
| Valine | 1.73E-01 (1.22E-01 - 2.47E-01) | 1.40E-01 (8.72E-02 - 1.87E-01) | 0.0075 |
| Creatinine | 5.82E-02 (4.17E-02 - 7.73E-02) | 5.06E-02 (3.49E-02 - 6.37E-02) | 0.0177 |
| **Female associated:** | | | |
| XL-HDL- Particles | 2.28E-07 (0 - 5.50E-07) | 3.87E-07 (0 - 7.01E-07) | 0.0165 |
| XL-HDL- total lipids | 2.27E-01 (0 - 5.61E-01) | 3.87E-01 (0 - 7.04E-01) | 0.0176 |
| XL-HDL- phospholipids | 1.24E-01 (0 - 2.77E-01) | 2.21E-01 (0 - 3.99E-01) | 0.0069 |
| XL-HDL- cholesterol | 9.43E-02 (0 - 2.78E-01) | 1.55E-01 (0 - 2.90E-01) | 0.0493 |
| XL-HDL- free cholesterol | 2.34E-02 (0 - 7.61E-02) | 4.25E-02 (0 - 8.50E-02) | 0.0278 |
| L-HDL- Particles | 8.66E-07 (0 - 1.59E-06) | 1.33E-06 (8.29E-07 - 2.34E-06) | 0.006 |
| L-HDL- total lipids | 5.44E-01 (0 - 1.01) | 8.39E-01 (5.15E-01 - 1.49) | 0.006 |
| L-HDL- phospholipids | 2.67E-01 (0 - 4.75E-01) | 4.00E-01 (2.74E-01 - 6.93E-01) | 0.0057 |
| L-HDL- cholesterol | 2.58E-01 (0 - 5.13E-01) | 4.11E-01 (2.18E-01 - 7.48E-01) | 0.0074 |
| L-HDL- cholesterol ester | 2.05E-01 (0 - 3.97E-01) | 3.22E-01 (1.73E-01 - 5.79E-01) | 0.0076 |
| L-HDL- free cholesterol | 5.26E-02 (0 - 1.16E-01) | 8.93E-02 (4.48E-02 - 1.70E-01) | 0.0067 |
| L-HDL- triglycerides | 1.84E-02 (0 - 4.45E-02) | 2.75E-02 (1.25E-02 - 4.71E-02) | 0.0118 |
| S-HDL- triglycerides | 4.82E-02 (2.54E-02 - 7.28E-02) | 3.78E-02 (2.32E-02 - 5.47E-02) | 0.0128 |
| HDL-Diameter | 9.78 (9.4 - 1.01E+1) | 1.00E+01 (9.65 - 1.03E+01) | 0.006 |
| HDL- cholesterol | 1.20 (7.48E-01 - 1.73) | 1.48 (1.10 - 2.13) | 0.0111 |
| HDL2- cholesterol | 7.48E-01 (3.35E-01 - 1.24) | 1.01 (6.65E-01 - 1.63) | 0.0114 |
| HDL3- cholesterol | 4.47E-01 (4.12E-01 - 4.90E-01) | 4.68E-01 (4.35E-01 - 5.10E-01) | 0.023 |
| ApoA1 | 1.33 (1.06 - 1.62) | 1.48 (1.25 - 1.85) | 0.027 |
| Sphingomyelin | 3.07E-01 (2.26E-01 - 4.27E-01) | 3.53E-01 (2.82E-01 - 4.28E-01) | 0.0339 |
| Docosahexaenoic acid | 1.01 (7.03E-01 - 1.45) | 1.22 (8.05E-01 - 1.78) | 0.0191 |
| Linoleic acid | 2.94E+01 (2.69E+1 - 3.39E+01) | 3.25E+01 (2.94E+01 - 3.64E+01) | 0.0003 |
| Omega-6 fatty acids | 3.43E+01 (3.06E+01 - 3.85E+01) | 3.73E+01 (3.32E+01 - 4.11E+01) | 0.0007 |
| Polyunsaturated fatty acids | 3.75E+01 (3.35E+01 - 4.23E+01) | 4.08E+01 (3.62E+01 - 4.43E+01) | 0.0005 |
| Citrate | 1.02E-01 (8.01E-02 - 1.30E-01) | 1.22E-01 (9.70E-02 - 1.51E-01) | 0.0007 |

**Supplemental Table 3: Metabolites significantly associated with adolescent healthy post-pubertal males and females.** All metabolites (Nightingale) with $p<0.05$ (unpaired t test) between males and females. P values were corrected for multiple testing by 10% false discovery rate (FDR) correction (Benjamini, Krieger and Yekutieli approach). P=Unpaired t-test. Key: brown (VLDL), green (HDL), grey (apolipoproteins), pink (fatty acids, %FA), yellow (other lipids), purple (amino acids), blue (glycolysis), black (other).

**Supplemental Table 4: Metabolites that overlap between gender group comparisons**

| Male vs Female **vs** Male GD | Male vs Female **vs** Female GD | Male GD **vs** Female GD | Total overlap between all comparisons |
|---|---|---|---|
| XL-HDL-P | L-HDL-P | L-HDL-P | L-HDL-P |
| XL-HDL-L | L-HDL-PL | L-HDL-L | L-HDL-PL |
| XL-HDL-PL | L-HDL-TG | L-HDL-PL | L-HDL-TG |
| XL-HDL-C | HDL-C | L-HDL-TG | HDL-C |
| XL-HDL-FC | HDL2-C | M-HDL-PL | HDL2-C |
| L-HDL-P | ApoA1 | HDL-C | ApoA1 |
| L-HDL-PL | DHA | HDL2-C | |
| L-HDL-C | | ApoA1 | |
| L-HDL-CE | | | |
| L-HDL-FC | | | |
| L-HDL-TG | | | |
| HDL-D | | | |
| HDL-C | | | |
| HDL2-C | | | |
| HDL3-C | | | |
| ApoA1 | | | |

**Supplemental Table 4: Metabolites that overlap between gender group comparisons**
Table displaying the overlapping metabolites (Venn diagram Figure 3G) that were significantly different between gender comparisons from Figure 3. GD; Gender dysphoria.

**Supplemental Table 5: Demographic and clinical comparison between JSLE males and females within the cohort**

| Demographic | Male | Female | P value |
|---|---|---|---|
| **Number** | 12 | 23 | - |
| **Age** | 18 (15-23) | 20 (14-25) | 0.39 |
| **White** | 3 (25%) | 10 (43%) | 0.46 |
| **Asian** | 6 (50%) | 7 (30%) | 0.29 |
| **Black** | 3 (25%) | 4 (17%) | 0.67 |
| **Other** | 0 (0%) | 2 (9%) | 0.54 |
| **Age of onset** | 13 (0-18) | 12 (5-18) | 0.32 |
| **Clinical feature** | | | |
| **SLEDAI (active disease>4)** | 2 (0-10) | 2 (0-10) | 0.62 |
| **Neurological involvement** | 1 (8%) | 1 (4%) | 1.00 |
| **Serositis involvement** | 2 (17%) | 6 (26%) | 0.31 |
| **Cutaneous involvement** | 10 (83%) | 21 (92%) | 0.69 |
| **Heamatological involvement** | 5 (42%) | 9 (39%) | 1.00 |
| **Musculoskeletal involvement** | 9 (75%) | 21 (91%) | 0.31 |
| **Renal involvement** | 4 (33%) | 5 (22%) | 0.69 |
| **RF_Result (positive)** | 1 (8%) | 2 (9%) | 1.00 |
| **LA_Result (positive)** | 2 (17%) | 2 (9%) | 0.59 |
| **ENA (positive)** | 5 (42%) | 14 (61%) | 0.31 |
| **ESR (NR=<20)** | 6.5 (2-38) | 13 (2-127) | 0.23 |
| **dsDNA_Result (NR=<50)** | 47 (2-2827) | 58 (1-17441) | 0.46 |
| **C3 (NR=0.9-1.8)** | 1 (0.33-1.36) | 1.11 (0.35-1.64) | 0.28 |
| **Lymphocyte count (NR=1.3-3.5)** | 1.49 (0.68-3.8) | 1.63 (0.59-3.08) | 0.78 |
| **Treatment** | | | |
| **Hydroxychloroquine** | 11 (92%) | 20 (87%) | 1.00 |
| **Mycophenolate mofetil** | 5 (42%) | 13 (57%) | 0.49 |
| **Prednisolone** | 5 (42%) | 12 (52%) | 0.72 |
| **Vitamin D** | 2 (17%) | 6 (26%) | 0.19 |
| **Methotrexate** | 1 (8%) | 2 (9%) | 1.00 |
| **Azathioprine** | 4 (33%) | 3 (13%) | 0.20 |

**Supplemental Table 5: Demographic and clinical comparison between JSLE males and females within the cohort**

SLEDAI score was calculated, a score greater than 6 represents active disease [155]. Other common clinical measures of disease are shown as well as treatments. Rituximab treatment was avoided in the cohort. Fisher's exact test* or unpaired t-test was used. Abbreviations: NR: Normal ranges, SLEDAI: Systemic Lupus Erythematosus Disease Activity Index, RF: Rheumatoid factor, LA: Lupus Anticoagulant, dsDNA: Anti-double-stranded-DNA antibodies, C3: Complement component 3, LC: Lymphocyte count.

### Supplemental Figures

Supplemental Figure 1

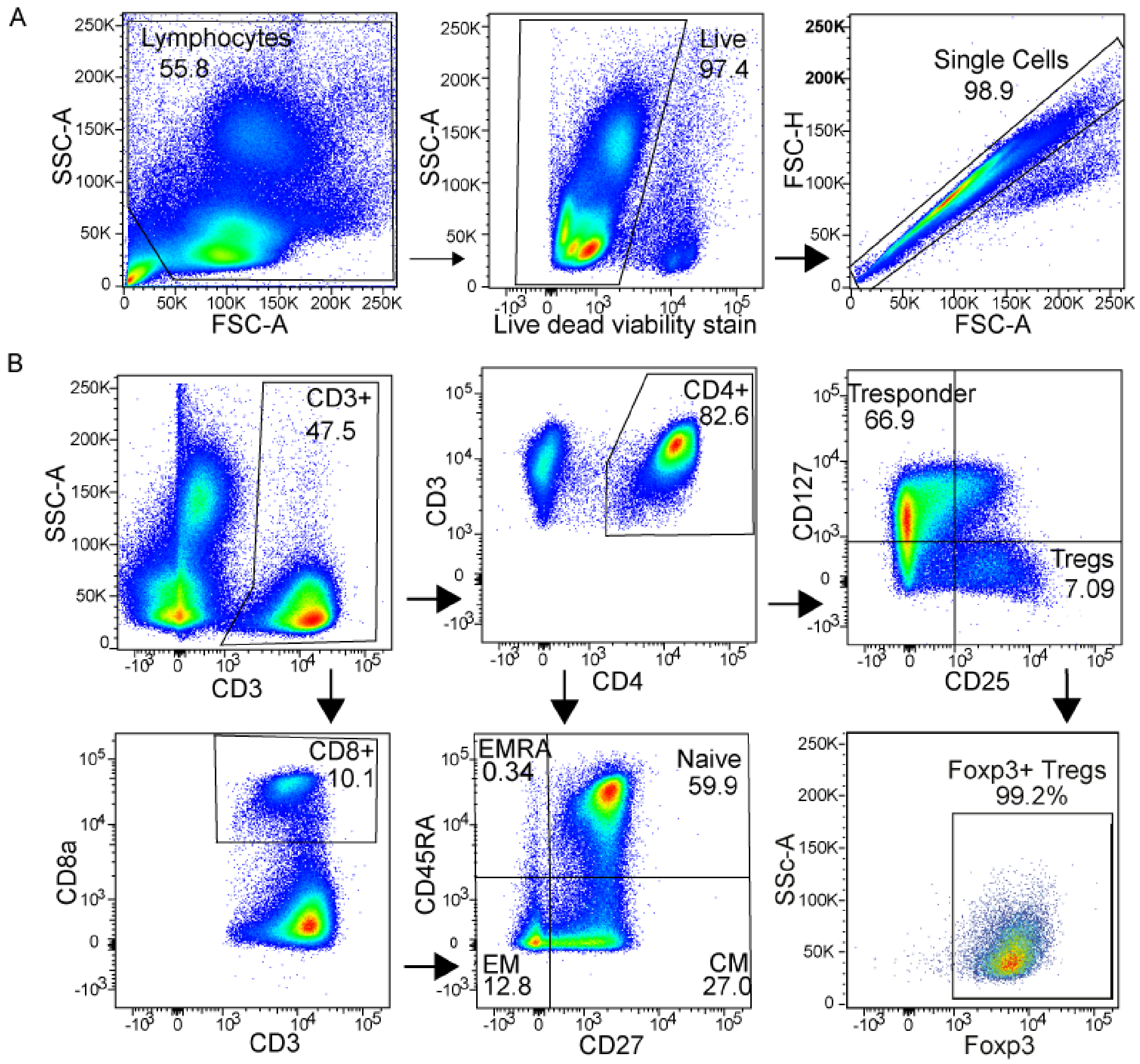


**Supplemental Figure 1: Gating strategies for flow cytometry immunophenotyping**
(A and B) Representative gating strategies from a healthy donor used to identify T-cell subsets. PBMC's were stained with antibodies outlined in methods. Samples were analysed using Fortessa X20 flow cytometer and Flowjo software. Labels represent the cell population within the gate. (B) Regulatory T-cells (Tregs), central memory (CM), effector memory (EM). Representative Foxp3 percentage expression is shown where PBMC's were stained separately with CD4, CD127, CD25 and intracellularly stained with Foxp3 to validate the purity of Tregs taken from a CD25+CD127- gate.

Supplemental Figure 2

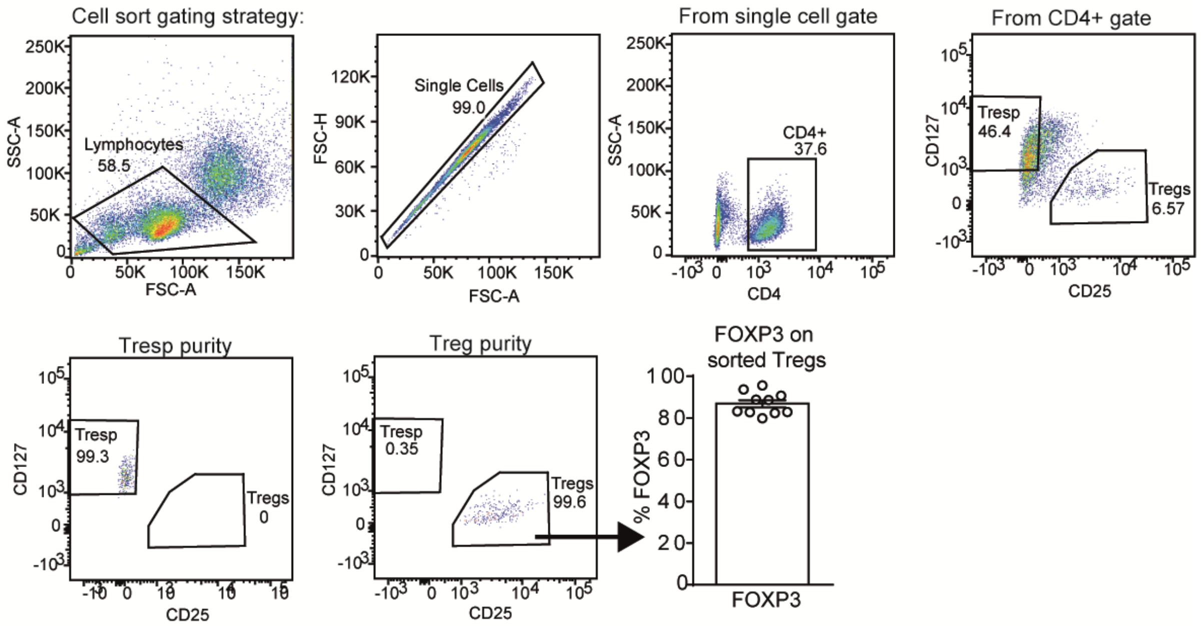


**Supplemental Figure 2: Gating strategy for Tregs and Tresp cell sorting and FOXP3 purity checks**

PBMCs were surface stained with CD4, CD25 and CD127 and sorted by a FACS Aria cell sorter. A small aliquot of Tregs were tested for purity each time through intracellular staining of FOXP3. Tregs and Tresp cells were then used for suppression assays or qPCR.

Supplemental Figure 3

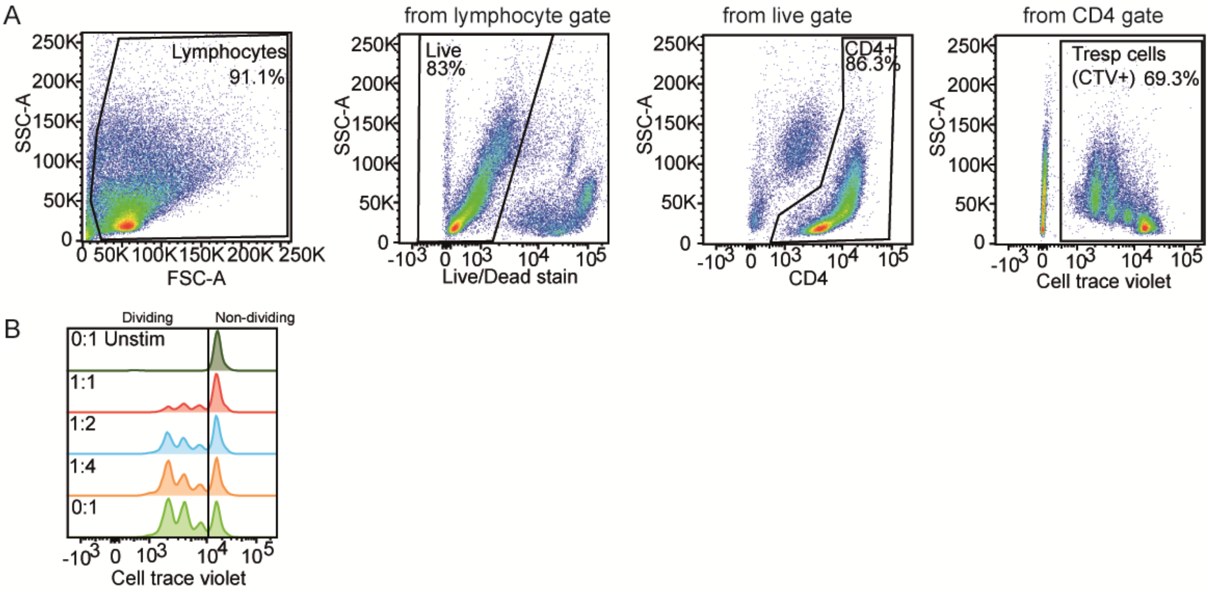


**Supplementary Figure 3: Gating strategy for Treg suppression assay analysis**
**(A)** Gating strategy for identification of Tresp (cell trace violet (CTV) positive) cells. **(B)** Representative plot of Tresp CTV expression through rounds of proliferation at varying ratios of Treg:Tresp cells. Non-dividing and dividing cells are labelled.

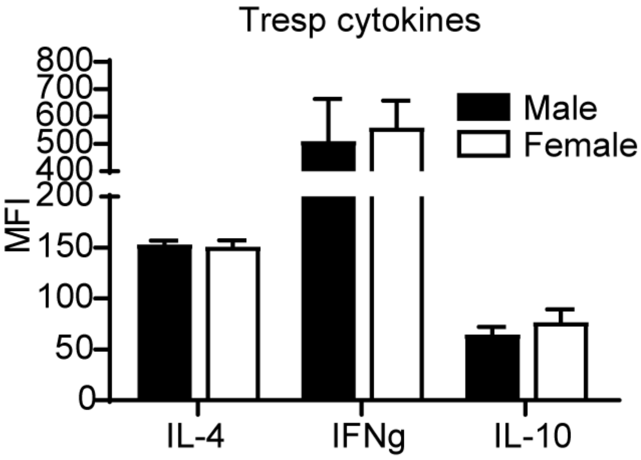


**Supplemental Figure 4: Cytokine and transcription factor expression on Tresp cells between healthy males and females**

Ex vivo PBMCs from 8 male and 12 female healthy donors were surface stained with CD4, CD25 and CD127 to detect Tresp cells and intracellularly for cytokines IL-4, IFNγ and IL-10 after 4hr stimulation with PMA/ionomycin/golgi plug. Samples were analysed using Fortessa X20 flow cytometer and Flowjo software. Mean+SE.

Supplemental Figure 5

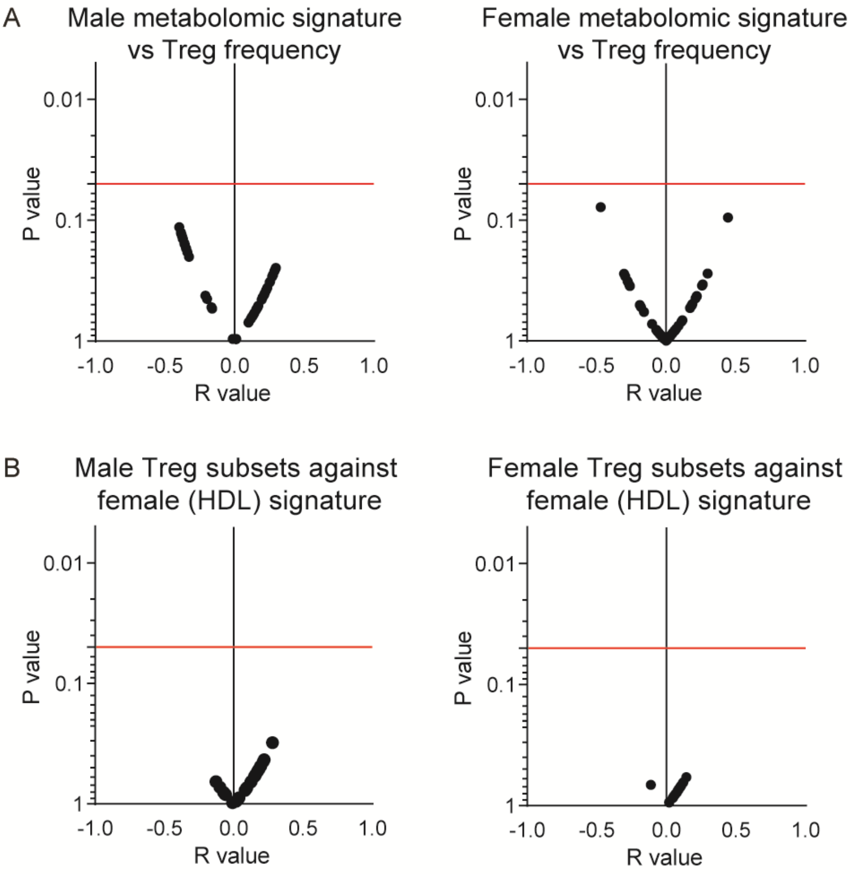


**Supplemental Figure 5: Correlations between male or female metabolomic signatures and Treg phenotypes (A)** Volcano plots showing the correlation (r value) and Log10 p values between the male (left) or female (right) metabolomic signature and the respective total Treg frequency. **(B)** Volcano plots showing the correlation (r value) and Log10 p values between the male (left) or female (right) Miyara et.al Treg functional subsets and the female (HDL dominated) metabolomic signature (Supplementary Table 3). Red line=p=0.05.

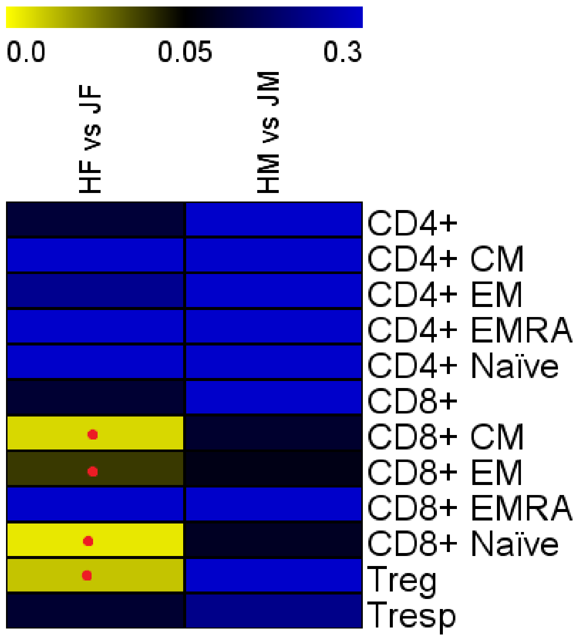


**Supplementary Figure 6: Comparison of immunophenotype between healthy and JSLE males and females**

Heatmap comparing T-cell subset immunophenotyping (gating strategy Supplementary Figure 1) of age and ethnically matched healthy/JSLE females (n=22/23) and males (n=17/12). Unpaired t-test. Red dot=significant p values following false discovery rate adjustment for multiple comparisons.
